# Genome-wide quantification of RNA flow across subcellular compartments reveals determinants of the mammalian transcript life cycle

**DOI:** 10.1101/2022.08.21.504696

**Authors:** Brendan M. Smalec, Robert Ietswaart, Karine Choquet, Erik McShane, Emma R. West, L. Stirling Churchman

## Abstract

Dissecting the myriad regulatory mechanisms controlling eukaryotic transcripts from production to degradation requires quantitative measurements of mRNA flow across the cell. We developed subcellular TimeLapse-seq to measure the rates at which RNAs are released from chromatin, exported from the nucleus, loaded onto polysomes, and degraded within the nucleus and cytoplasm. These rates varied substantially, yet transcripts from genes with related functions or targeted by the same transcription factors and RNA binding proteins flowed across subcellular compartments with similar kinetics. Verifying these associations uncovered roles for DDX3X and PABPC4 in nuclear export. For hundreds of genes, most transcripts were degraded within the nucleus, while the remaining molecules were exported and persisted with stable lifespans. Transcripts residing on chromatin for longer had extended poly(A) tails, whereas the reverse was observed for cytoplasmic mRNAs. Finally, a machine learning model identified additional molecular features that underlie the diverse life cycles of mammalian mRNAs.

## Introduction

The life cycles of mRNAs are dynamic and diverse. Thousands of mRNAs are produced per minute in a typical mammalian cell (Carter et al., 2005; Schofield et al., 2018). Before they can be translated, mRNAs must flow across subcellular compartments, including release from chromatin and export from the nucleus. In the cytoplasm, ribosomes are loaded onto mRNAs and transcripts are ultimately degraded. These transitions between compartments are controlled by numerous regulatory mechanisms. Accordingly, not all mRNAs are destined for this stereotypical trajectory, and those that are, do not flow across the cell at the same rates. Thus, RNA flow impacts cell function by determining the dynamic pool of mRNAs available for translation.

On chromatin, mRNAs are synthesized by RNA polymerase II and undergo extensive processing, including splicing and polyadenylation. The time required to excise introns from pre-mRNAs varies significantly, ranging from seconds to tens of minutes (Drexler et al., 2020; Martin et al., 2013; Pai et al., 2017; Rabani et al., 2014; Reimer et al., 2021; Wachutka et al., 2019; Wan et al., 2021). In some cases, splicing regulates the nuclear dynamics of nascent RNA (Mauger et al., 2016; Ninomiya et al., 2011; Pandya-Jones et al., 2013; Yeom et al., 2021). In the nucleus, transcripts are subjected to either degradation or export. Nuclear degradation targets improperly processed mRNAs (Bresson et al., 2015; Davidson et al., 2012; Meola et al., 2016; Pendleton et al., 2018) but also serves additional regulatory roles for specific transcripts (*Gudipati et al., 2012*). By contrast, some loci are tethered near the nuclear pore complex to promote rapid export of transcripts (Blobel, 1985; Rohner et al., 2013; Scholz et al., 2019).

In the cytoplasm, ribosomes are loaded onto transcripts with different kinetics, partially influenced by 5’UTR length and structure (Lai et al., 2008; Leppek et al., 2018; Parsyan et al., 2009; Pisareva et al., 2008; Soto-Rifo et al., 2012), and promoter elements (Zid and O’Shea, 2014). Finally, mRNAs undergo degradation, a process that can be driven by both poly(A) tail deadenylation and targeting by microRNAs (Bartel, 2018; Eisen et al., 2020a, 2020b; Passmore and Coller, 2022). Each of these processes vary in duration across genes and impact the subcellular fates of transcripts, either directly or indirectly through feedback loops. In total, RNA half-lives vary greater than 100-fold between different protein-coding transcripts (Dölken et al., 2008; Friedel et al., 2009; Herzog et al., 2017; Rabani et al., 2011; Schofield et al., 2018; Schwanhäusser et al., 2011). However, half-lives measured at the whole-cell level only measure the time between synthesis and decay, obscuring the dynamics of mRNA transitions across subcellular compartments.

Around 50 years ago, metabolic labeling in mammalian cells with radiolabeled nucleotide precursors shed light on bulk RNA flow and metabolism, but these experiments could not resolve transcript-specific behaviors (Darnell et al., 1973). Several methods have been developed to assay the rates of RNA flow for one or a few transcripts. For example, single-molecule microscopy approaches track reporter RNAs throughout mammalian cells (Halstead et al., 2015; Hoek et al., 2019; Mor et al., 2010; Shav-Tal et al., 2004). Endogenous RNAs have been studied using single-molecule RNA FISH combined with mathematical modeling, yielding nuclear and cytoplasmic RNA half-lives of a handful of genes in mouse tissue (Bahar Halpern et al., 2015) and allowing for the quantification of the entire life cycle of an individual transcript in *Arabidopsis* (Ietswaart et al., 2017; Wu et al., 2016). To determine the rates at which RNAs flow across compartments with higher throughput, induced genes have been monitored over time and modeled to estimate subcellular turnover (Battich et al., 2015; Bhatt et al., 2012; Rabani et al., 2014), yet the kinetics observed for these transcripts may not extend to all genes or to cellular contexts that do not involve gene induction. Recently, metabolic labeling has been used to globally study specific stages of RNA lifespans (Berry et al., 2022; Chen and van Steensel, 2017; Schott et al., 2021), but have not comprehensively characterized the entire life cycle of an mRNA across multiple compartments in mammalian cells, and none have investigated degradation in the nucleus.

Here, we quantify the rates of mRNA flow across mammalian cells genome-wide. We start by introducing subcellular TimeLapse-seq, a method that measures RNA turnover with subcellular resolution, and couple this technique with kinetic modeling to estimate the rates at which mRNAs flow across subcellular compartments. We measured RNA half-lives on chromatin, in the nucleus, and in the cytoplasm, and additionally measured nuclear export and polysome loading rates for all expressed genes in mouse NIH-3T3 and human K562 cells. Strikingly, for ~5–10% of genes, mRNA flow was predicted to involve substantial nuclear degradation. We found that RNA flow rates varied widely (>100-fold) between different genes and subcellular compartments. Our results demonstrate that functionally related genes undergo similar rates of RNA flow. The targets of many RNA binding proteins (RBPs) exhibit different RNA flow rates compared to other genes, and these differences dissipated upon RBP perturbation. Measurement of poly(A) tails with subcellular resolution revealed that tail lengths reflect subcellular RNA half-lives. Finally, we identified the strongest genetic and molecular features that are predicted to determine RNA flow through machine learning, including transcription factors and sequence elements. Collectively, our findings provide a comprehensive characterization of the dynamics of an mRNA throughout its life cycle within a mammalian cell.

## Results

### Subcellular TimeLapse-seq measures the fraction of newly synthesized RNA across subcellular compartments

To quantity RNA turnover genome-wide with subcellular resolution, we developed subcellular Timelapse-seq, a method that combined metabolic labeling with the biochemical purification of distinct RNA populations from different cellular compartments (Figure 1A). To generate the samples for TimeLapse-seq, we pulse labeled human K562 and mouse NIH-3T3 cells with 4-thiouridine (4sU) for 0, 15, 30, 60, and 120 minutes. Because high concentrations of 4sU can broadly impact gene expression (Burger et al., 2013), we minimized the concentration and duration of 4sU exposure and confirmed that the addition of 4sU did not affect mRNA subcellular localization (Figure S1A-B) and led to minimal disruption of gene expression levels within each subcellular compartment (Figure S1C). Following each 4sU pulse, we biochemically purified chromatin-associated, nuclear, and cytoplasmic RNA (Figure 1B) (Mayer and Churchman, 2017), as well as polysome-bound RNA (Figure S1D). We also collected total cellular RNA. We then quantified the amount of newly synthesized RNA in each sample by performing TimeLapse-seq, a nucleotide conversion protocol that detects 4sU-labeled RNA within a mixture of labeled and unlabeled RNAs based on the presence of 4sU-induced T>C sequencing mismatches (Figure 1A) (Schofield et al., 2018).

**Figure 1:**
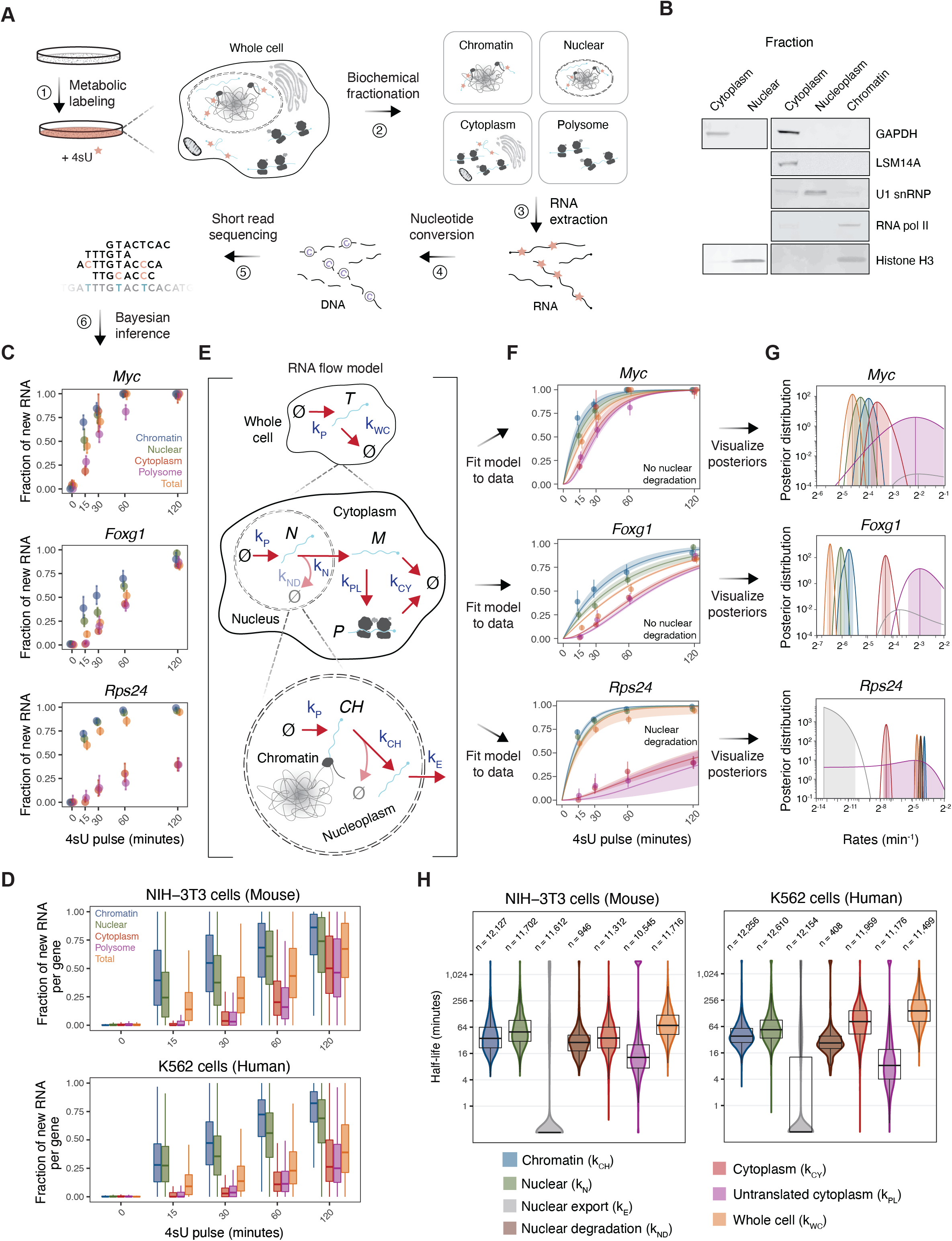
Subcellular TimeLapse-seq and kinetic modeling estimate genome-wide RNA flow rates. (A) Schematic representing subcellular TimeLapse-seq. Cells were pulse-labeled with 4-thiouridine (4sU) and biochemically fractionated into subcellular compartments. TimeLapse-seq libraries were prepared from each sample and the fraction of newly synthesized RNA per gene was estimated (see Fig. S2B and Methods for details). (B) Western blot of subcellular marker proteins: GAPDH and LSM14A, cytoplasmic proteins; U1 snRNA, nucleoplasmic protein; histone H3 and RNA pol II, chromatin proteins. (C) Subcellular TimeLapse-seq data for example genes (*Myc, Foxg1, Rps24*) in mouse NIH-3T3 cells. Dots represent the fraction of new RNA MAP values for one replicate, while vertical lines represent the 95% credible intervals (CIs). (D) Genome-wide subcellular TimeLapse-seq data for all protein-coding genes in human K562 and mouse NIH-3T3 cells. Fraction of new RNA MAP values for each gene are shown for one replicate. (E) Schematic of the RNA flow model (see Fig. S3A and Methods for details). (F) RNA flow model fit to subcellular TimeLapse-seq data for the example genes shown in (C). The dark lines represent the RNA flow rate MAPs while the ribbons show the 95% CIs. Colors are consistent with the RNA populations in (C). (G) Posterior distributions for each RNA flow rate modeled in (F) with the MAPs represented with vertical lines and 95% CIs in shading. Colors are consistent with the rates shown below in (H). (H) Genome-wide subcellular half-lives for all protein-coding genes in mouse NIH-3T3 and human K562 cells. Nuclear degradation rates are only included for genes best explained by this model. Mean half-lives are shown and the number of genes noted.

Previous applications of nucleotide conversion approaches have used labeling conditions that achieved high 4sU incorporation rates, aiding the identification of labeled RNAs (Erhard et al., 2019; Herzog et al., 2017; Schofield et al., 2018). Therefore, the lower 4sU incorporation rates inherent to our minimal labeling conditions necessitated a new approach to estimate the fraction of newly synthesized RNA (Figure S2A-B). We developed a binomial mixture model to estimate upper and lower bounds on the T>C conversion rates for each 4sU pulse time and compartment (Figure S2B). We then inputted these conversion rates into GRAND-SLAM (Jürges et al., 2018) and combined the outputs to quantify the posterior distribution on the fraction of newly synthesized RNA per gene within each sample (Figure S2B). We validated our approach with a NanoStrings-based assay that does not rely on sequencing and predictions of T>C conversions (Figure S2C). With this technique, we observed fractions of new nuclear, cytoplasm, and total RNA similar to those estimated by subcellular TimeLapse-seq for select genes with both fast and slow turnover (Figure S2D-E), validating the robustness of our analysis pipeline across a range of mismatch rates.

Genome-wide, we observed an increase in the proportion of new RNA with increasing pulse durations within each compartment (Figure 1C-D). Furthermore, at each time point, we saw delays in the fraction of new RNA across chromatin, nuclear, cytoplasm, and polysome fractions (Figure 1C-D), even for genes with relatively fast turnover (e.g., *Myc*, Figure 1C). This result confirmed that our assay has the time resolution suitable for detecting RNA flow across subcellular compartments.

### Kinetic modeling of RNA flow across subcellular compartments

To estimate the rates at which RNAs flow across subcellular compartments for each gene, we fit a kinetic model consisting of a system of ordinary differential equations to our subcellular TimeLapse-seq data (Figure 1E). By coupling this model to the Bayesian inference framework of GRAND-SLAM (Figure S3A), we estimated the Bayesian posterior probability distribution of each flow rate per gene (Figure 1F-G, Table S1). Using the posterior mean rate (*k*), half-lives are then calculated as 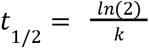. We determined the half-lives of RNAs on chromatin (“chromatin half-lives”), in the nucleus (“nuclear half-lives”), and in the cytoplasm (“cytoplasm half-lives”). We estimated rates of nuclear export (yielding “nuclear export half-lives”) and the rates at which polysomes are loaded onto RNAs after entry into the cytoplasm (yielding “untranslated cytoplasm half-lives”). Finally, we estimated the rates at which RNAs are turned over at the whole-cell level (yielding “whole-cell half-lives”), using only the total RNA data (Figure 1E).

This initial model fit the data for most genes well (e.g. *Myc* and *Foxg1*, Figure 1F, Table S1). The model predicts that whole-cell half-lives represent the sum of the nuclear and cytoplasmic half-lives, because it did not include nuclear degradation of mRNA (Schmid and Jensen, 2018) (Figure 1E). However, in the presence of substantial nuclear RNA degradation, this relation will no longer hold true, as many RNAs never exist in the cytoplasm and whole-cell data will no longer be fit by the model. We noticed this trend for a small group of genes in both cell lines, including *Rps24* in NIH-3T3 (Figure S3B). Therefore, we extended our model by including a nuclear RNA degradation rate. To determine if adding this parameter yields a better fit for each gene, we calculated a Bayes factor (Kass and Raftery, 1995), namely, the ratio of likelihoods between the nuclear degradation model (alternative hypothesis) and the model with no nuclear degradation (null hypothesis). In the absence of nuclear degradation, we were not able to predict the total RNA data for *Rps24* (Figure S3B), whereas a model with nuclear degradation was successful (Figure 1F, Bayes factor=10^136^).

Chromatin, nuclear, nuclear export, nuclear degradation, cytoplasm, and whole-cell half-lives were strongly correlated between biological replicates, with small 95% credible intervals (CIs) (Figure S3D,E, Table S1, Pearson correlation r>0.75), and strong correspondence with previously reported whole-cell half-lives (Figure S3C, Pearson correlation r=0.65). However, most genes had larger 95% CIs for their untranslated cytoplasm half-lives than for other flow rates, and the half-lives also exhibited greater variation between biological replicates (Figure S3D,E, r=0.44 in K562 and r=0.45 in NIH-3T3). Nevertheless, the 95% CIs reproduced between replicates (overlap for 86% genes in NIH-3T3, 79% genes in K562) and did not exceed the variation between genes. To assess the robustness of our RNA flow rates across modeling approaches, we compared our Bayesian distributions to least-squares estimates. Although chromatin, nuclear, cytoplasm, and whole-cell half-lives were similar (r>0.67), the least squares model failed to estimate untranslated cytoplasm half-lives (r=0.42 in K562, r=0.17 in NIH-3T3) due to its inability to take measurement uncertainties into account (Figure S3E). We conclude that our Bayesian approach is capable of robustly quantifying the flow of RNAs across subcellular compartments for endogenously expressed genes in mouse and human cells.

### Wide gene-to-gene variability in RNA flow rates

In both cell lines, RNA flow rates varied considerably between genes (Figure 1H). For 90% of genes, chromatin half-lives ranged from 19 to 120 minutes in K562 and from 14 to 200 minutes in NIH-3T3, with medians around 40 minutes. Nuclear RNA half-lives in both cell lines were often somewhat longer than, but highly correlated with, chromatin half-lives (r>=0.75, Figure S4A-B), with median nuclear half-lives around 45 minutes that ranged between 20 and greater than 260 minutes for 90% of genes. As expected from the similarity between chromatin and nuclear half-lives for most genes, more than 70% of all genes had very rapid nuclear export half-lives of less than 10 minutes. Thus, most mRNAs within the nucleus are associated with chromatin (Figure S4C). Nevertheless, slow export does occur; at least 10% of genes had longer export half-lives (>30 minutes) in both K562 and NIH-3T3.

In the cytoplasm, mRNAs were more stable in K562 cells with a median half-life of 78 minutes, compared to 36 minutes in NIH-3T3. Thus, turnover of mRNAs at whole-cell resolution was also slower in K562 cells than in NIH-3T3 cells, with median whole-cell half-lives of 140 and 71 minutes, respectively. In both cell lines, cytoplasm half-lives varied greatly, ranging from 10 to 140 minutes for 90% of genes in NIH-3T3 and from 20 to 260 minutes in K562 cells. Notably, differences in cytoplasm mRNA stability did not correspond with differences in rates of polysome loading, which occurs relatively quickly after nuclear export. In both cell lines, the median untranslated cytoplasm half-life was less than 15 minutes, with more than 75% of genes having half-lives less than 30 minutes. Untranslated cytoplasm half-lives and cytoplasm half-lives were uncorrelated in K562 (r=0.06) and weakly correlated in NIH-3T3 (r=0.28) (Figure S4D-E). Thus, loading of RNAs onto polysomes is not strongly coupled with cytoplasmic RNA turnover. On average, mRNAs for most genes spent longer in the nucleus than the cytoplasm in NIH-3T3 cells. Although, in both cell lines, nuclear and whole-cell half-lives (r=0.66 in K562, r=0.83 in NIH-3T3) were more strongly correlated than cytoplasm and whole-cell half-lives (r=0.36 in K562, r=-0.06 in NIH-3T3) (Figure S4F-G). As expected, we accurately estimated the whole-cell half-life by adding the nuclear and cytoplasm half-lives for most genes, but could not do so for genes predicted to undergo nuclear RNA degradation (Figure S4H-I).

### Genes encoding transcripts <u>p</u>redicted to <u>u</u>ndergo <u>n</u>uclear <u>d</u>egradation (PUNDs) are conserved across mouse and human cells

The Bayes factors for 9% (n= 946/11,109) of genes in NIH-3T3 cells and 4% (n= 408/10,971) of genes in K562 cells reproducibly exceeded 100 (Figure 2A, S5A, Table S2), indicating that the model with nuclear degradation is at least 100 times more likely to explain the subcellular TimeLapse-seq data than the model without nuclear degradation, thus providing decisive evidence (Kass and Raftery, 1995) that transcripts of these genes undergo nuclear degradation. Hereafter, we refer to these genes as PUNDs (<u>p</u>redicted to <u>u</u>ndergo <u>n</u>uclear <u>d</u>egradation). The model predicts that the large majority (>85%) of transcripts produced by PUNDs are degraded in the nucleus.

**Figure 2:**
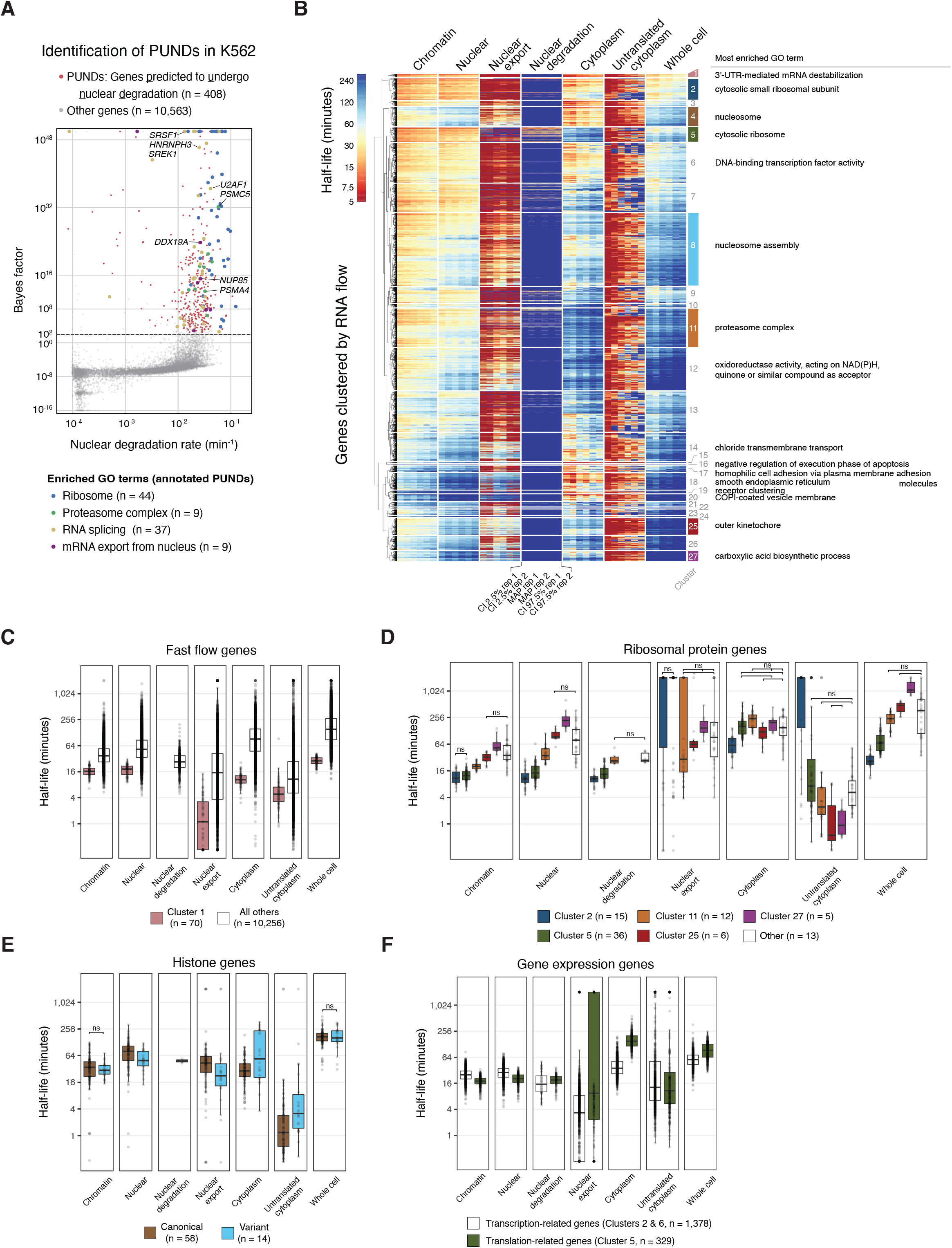
Genes with similar rates of RNA flow across the cell are functionally related. (A) Genes with a Bayes factor >100 in both replicates were labeled as those <u>p</u>redicted to <u>u</u>ndergo <u>n</u>uclear <u>d</u>egradation (PUNDs, shown in red) in human K562 cells. PUNDs are additionally colored by their associated enriched GO terms. (B) Hierarchical clustering of human genes according to their RNA flow rates (MAPs and 95% credible intervals). The most enriched GO annotations for each cluster are displayed on the right (full results in Table S3). (C) Fast flow genes, i.e. those in cluster 1 in (B), were enriched for functions related to intracellular signaling and response to stimuli. All comparisons were statistically significant (p<0.0001, Wilcoxon test). (D) Half-lives of ribosomal protein genes (RPGs) from each cluster where they were enriched (clusters 2, 5, 11, 25, and 27 in (B)). The number of RPGs within each cluster is noted. Non-significant comparisons are indicated as “ns” and all other comparisons were statistically significant (p<0.05, Wilcoxon test). (E) Half-lives of histone genes. Canonical, replication-dependent histone genes were enriched in cluster 4, while non-canonical, replication-independent histone genes (including histone variants) were enriched in cluster 8. The number of histone genes of each type is noted, and significance was noted as in (D). (F) Half-lives of clusters 2 and 6, containing genes related to transcription, and cluster 5, containing genes related to translation. The number of genes in each group is noted, and significance was noted as in (D).

We performed Gene Ontology (GO) enrichment analysis on our PUND gene lists (Table S3) and found many common enriched terms between K562 and NIH-3T3 PUNDs, including those pertaining to the ribosome, RNA splicing, and nuclear mRNA export (Figure 2A, S5A). Ribosomal protein genes were highly enriched in PUNDs in both cell lines, including 24 individual ribosomal protein gene homologs. In total, we found 133 homologous genes that were shared between both cell lines, a significant overlap (Fisher’s exact test: p=2.3×10^−14^) (Figure S5B). Given the common functions of PUNDs, as well as the overlap of individual PUNDs between cell lines, we conclude that nuclear degradation is a conserved regulatory feature that acts on select transcripts.

### Functionally related genes exhibit similar RNA flow across subcellular compartments

Genes with related functions tend to be co-regulated such that their mRNAs have similar rates of whole-cell turnover (Dölken et al., 2008; Friedel et al., 2009; Herzog et al., 2017; Rabani et al., 2011; Schofield et al., 2018; Schwanhäusser et al., 2011). To determine whether subcellular RNA flow rates may likewise serve a regulatory role, we first performed hierarchical clustering on all genes in human K562 cells based on their subcellular half-lives (Figure 2B). By allowing for sufficient granularity, we identified 27 total clusters ranging from the order of 10 to 10^3^ genes with reproducibly distinct transcript kinetics (Figure 2B). We then performed GO enrichment analysis on each cluster and found that a majority have several enriched terms (Figure 2B, Table S3). Cluster 8, which contains the most genes (n=1,662), represents “canonical” RNA flow: median whole-cell half-lives of 150 min., long cytoplasmic and relatively short chromatin and nuclear residence, fast polysome loading, and no evidence for nuclear degradation (Figure 2B). However, the vast majority of genes (n=8,662) were divided into smaller groups that deviated from these canonical kinetics (Figure 2B). Genes involved in signal transduction and response to stimuli (e.g., *MYC, JUN*, and *CXCL2*) were overrepresented in a cluster of 70 genes with “fast flow” kinetics across all compartments (median whole-cell half-life 29 min.) (Figure 2C); however, even these genes spent more than half of their life cycle on chromatin (median half-life 17 min.). We corroborated this result independent of our clustering analysis by performing gene set enrichment analysis (GSEA) (Liberzon et al., 2015; Subramanian et al., 2005) of all genes ranked by each subcellular half-life and found that many hallmark gene sets involved in signaling pathways had significantly faster RNA flow across all compartments (e.g., TNF-alpha signaling, Figure S5C, Table S4). Thus, mRNAs with short half-lives across all compartments were enriched for signaling and sensing functions.

Ribosomal protein genes (RPGs) were enriched in only five clusters (Figure 2D, Table S2), rather than being uniformly distributed throughout all clusters (χ^2^ test p<10^−16^). Although all five of these clusters shared long cytoplasm half-lives, consistent with previous reports (Eisen et al., 2020a; Herzog et al., 2017; Munchel et al., 2011), other RNA flow rates differed. A majority of RPGs were PUNDs with short half-lives on chromatin, with either slow (cluster 2) or average (cluster 5) polysome loading kinetics. On the other hand, the remaining RPGs exhibited slower and more canonical RNA flow without nuclear degradation and fast polysome loading kinetics (clusters 11, 25, and 27). Thus, although not all RPGs exhibited the same rates of RNA flow, we nonetheless observed two distinct patterns of RNA flow that they tended to follow.

Histone genes were primarily enriched in two separate clusters (Figure 2B, clusters 4 and 8). Closer inspection revealed that the first group contained mostly canonical, replication-dependent histone genes, and the second contained variant histone genes (Figure 2E, Table S2). Whole-cell half-lives did not differ between these two groups; however, canonical histones had ~2-fold longer nuclear (median 79 min.) than cytoplasm (median 28 min.) half-lives (Figure 2E). Additionally, canonical histones were loaded onto polysomes very quickly, often within just a few minutes, consistent with a previous study (Schott et al., 2021). Thus, canonical histones experience relatively unique RNA flow, whereas variant histones behave more like the average protein-coding transcript.

Finally, genes involved in gene expression clustered into several distinct groups (Figure 2F). Most of the enriched GO terms in clusters 2 and 6 are related to transcription, including terms pertaining to transcription factors, RNA metabolism, and RNA polymerase II activity (Table S3). These genes included several mediator subunits, splicing factors, chromatin remodelers, and transcription factors (Table S1). By contrast, genes in cluster 5 are enriched for functions related to cytoplasmic translation, peptide biosynthesis, and ribosomal subunit biogenesis (Table S3), and include many translation factors (Table S1). Overall, genes involved in transcription had shorter whole-cell half-lives than genes involved in translation, consistent with multiple reports (Herzog et al., 2017; Schofield et al., 2018; Schwanhäusser et al., 2011; Yang et al., 2003) (Figure 2F). Surprisingly, despite their shorter whole-cell half-lives, genes involved in transcription had longer chromatin and nuclear half-lives and shorter cytoplasm half-lives compared to genes involved in translation (Figure 2F). Thus, genes involved in transcription and translation are enriched in clusters representing opposite RNA flow patterns.

### RNA flow rates are associated with RNA binding proteins

mRNAs exist as part of ribonucleoprotein complexes containing many RBPs, which control RNA metabolism and are likely regulators of RNA flow. To determine which RBPs correspond with specific RNA flow rates, we analyzed ENCODE eCLIP datasets for 120 RBPs in K562 cells (Van Nostrand et al., 2020), classified the mRNAs targets of each RBP, and identified all RBPs with target mRNAs that had significantly shorter or longer subcellular half-lives than non-targets (Figure 3A-B, Figure S6A-E, Table S5). We identified significant differences in the subcellular half-lives for nearly all (38/43) of the RBPs with fast or slow whole-cell half-lives (Figure 3B), allowing us to pinpoint exactly where in the cell each RBP may regulate RNA flow. Additionally, we identified significant differences in subcellular half-lives for the targets of an additional 37 RBPs that did not show differences at the whole-cell level, and a majority (25/37) of these RBPs only exhibited significant differences in chromatin and/or nuclear half-lives (Figure 3B).

**Figure 3:**
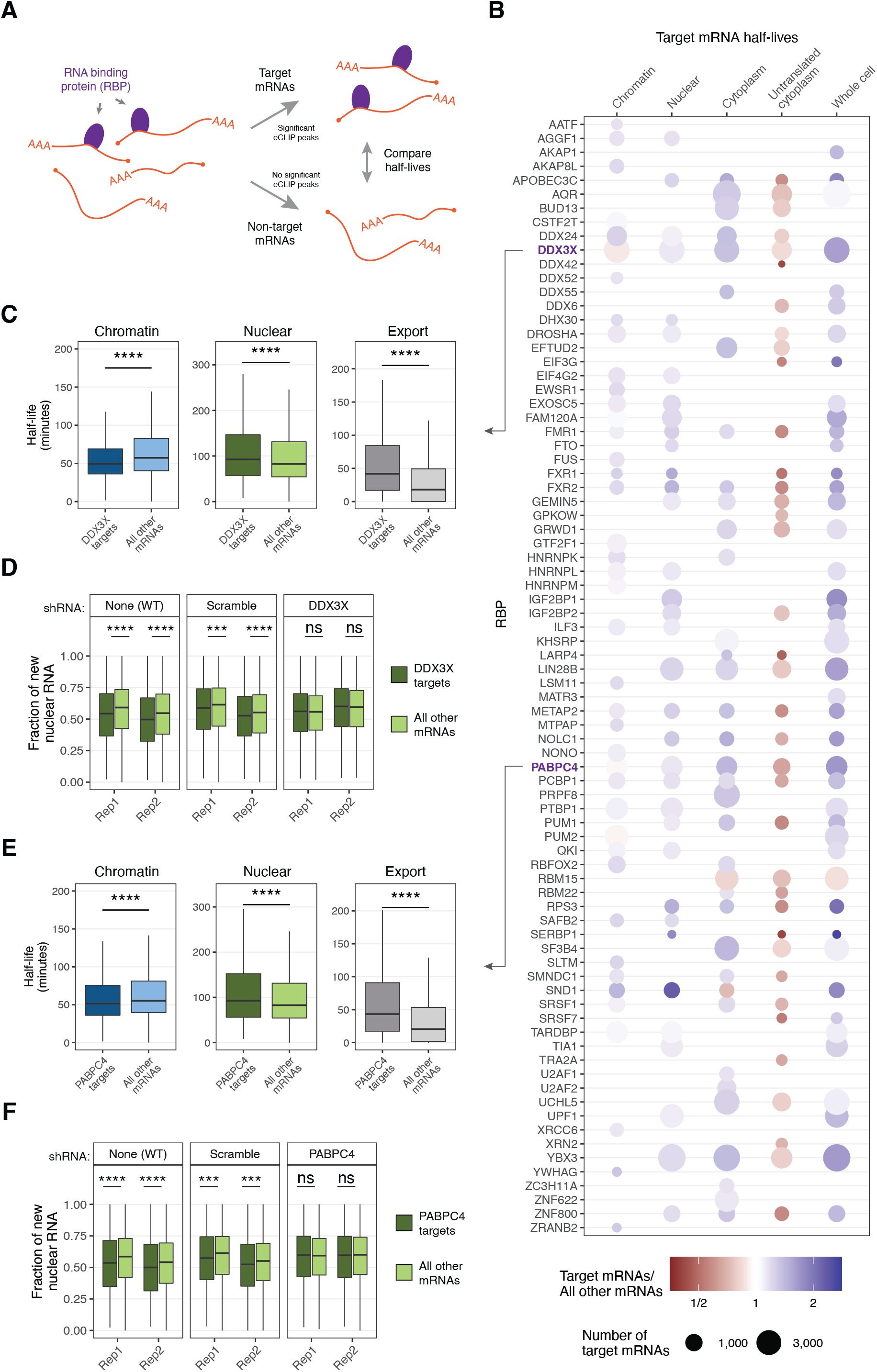
The targets of many RNA binding proteins (RBPs) exhibit distinctive RNA flow rates across the cell. (A) Schematic for RBP analysis. The mRNA binding targets of 120 RBPs in K562 were determined by identifying genes with significant eCLIP peaks published in (Van Nostrand et al., 2020). (B) All RBPs with targets that exhibited significantly fast or slow half-lives for target RNAs compared to non-target RNAs in both biological replicates (adjusted p<0.01, Wilcoxon test, Bonferroni multiple testing correction) across any RNA flow rate. The size of the dot indicates the number of target mRNAs with measured half-lives within each compartment and the color reflects the difference (red, faster; blue, slower) of the median target over non-target half-lives. (C) Half-lives of DDX3X mRNA targets and non-targets. The chromatin, nuclear, and nuclear export half-lives of targets are compared to non-target mRNAs (****: p<0.0001, ***: p<0.001, “ns:” not significant, Wilcoxon test). (D) Fraction of new nuclear RNA measured by subcellular TimeLapse-seq of DDX3X target mRNAs compared to all other mRNAs in wild-type cells, cells expressing a DDX3X-targeting shRNA, and cells expressing a scrambled shRNA. Significance was noted as in (C). Two biological replicates (“rep”) are shown. (E) Same as (C) for PABPC4 target genes. (F) Same as (D) for PABPC4.

### DDX3X and PABPC4 regulate nuclear export of target mRNAs

We next investigated the roles of RBPs in dictating RNA flow. We began by focusing on DDX3X, an RNA helicase with many roles in RNA metabolism (Kanai et al., 2004; Lai et al., 2008; Samir et al., 2019; Shih et al., 2012; Soto-Rifo et al., 2012; Yedavalli et al., 2004). DDX3X targets exhibited short chromatin half-lives and long nuclear half-lives, indicating slow export from the nucleus (Figure 3C, Figure S6A-B). To analyze the role of DDX3X in determining the flow rates of its target mRNAs, we used shRNAs to deplete DDX3X (Figure S6F) and performed subcellular TimeLapse-seq. Upon depletion, nuclear RNA turnover of DDX3X target mRNAs was no longer slower than non-targets (Figure 3D). Furthermore, DDX3X targets had long cytoplasm and whole-cell half-lives in wild-type cells (Figure 3B, Figures S6C,E), consistent with the association of DDX3X with several cytoplasmic RNA granules (Kanai et al., 2004; Samir et al., 2019; Shih et al., 2012). However, RNA turnover on chromatin, in the cytoplasm, and at the whole-cell level were not affected in the DDX3X knockdown (Figure S6H). We conclude that DDX3X regulates RNA flow at the step of nuclear export.

We observed similar RNA flow rates for targets of cytoplasmic poly(A) binding protein PABPC4, which had long cytoplasm and whole-cell half-lives (Figure 3B, Figures S6C,E), consistent with the function of this RBP in stabilizing transcripts with short poly(A) tails containing AU-rich motifs (Kini et al., 2014). However, PABPC4 targets had shorter chromatin half-lives and longer nuclear half-lives than other transcripts (Figure 3E). As before, we used shRNAs to deplete PABPC4 in K562 cells (Figure S6G) and performed subcellular TimeLapse-seq. Surprisingly, we detected no differences in the chromatin, cytoplasm, or whole-cell half-lives of target mRNAs upon PABPC4 depletion (Figure S6I). However, the target mRNAs no longer had longer nuclear half-lives (Figure 3F), implicating PABPC4 in nuclear export. Based on these findings, we conclude that although both PABPC4 and DDX3X bind to their mRNA targets throughout the cell, their depletion only affects RNA flow at the step of nuclear export. These observations highlight the ability of subcellular TimeLapse-seq to study the role of RBPs within different subcellular compartments and demonstrate that RBPs regulate RNA flow.

### mRNA poly(A) tail lengths are dynamic across and within subcellular compartments

Poly(A) tail lengths are connected to mRNA metabolism (Passmore and Coller, 2022), so we next analyzed the relationship between poly(A) tail length and RNA flow rates. We measured the tail length of each mRNA in chromatin, cytoplasm, and polysome fractions, as well as in total cellular RNA, using nanopore direct RNA sequencing (Figure 4A) (Workman et al., 2019). We confirmed that the 3’-end of >80% of sequenced RNAs mapped to annotated poly(A) sites (Figure S7A). To control for technical variations between sequencing runs, we included a set of six spike-in RNAs, each with a different poly(A) tail length. We calculated a poly(A) tail length size factor for each sample and used it to normalize endogenous RNAs poly(A) tail lengths (Figure S7B-C, methods). Within each fraction, the median tail length per gene correlated well between the two biological replicates (Figure S7D, Table S6).

**Figure 4:**
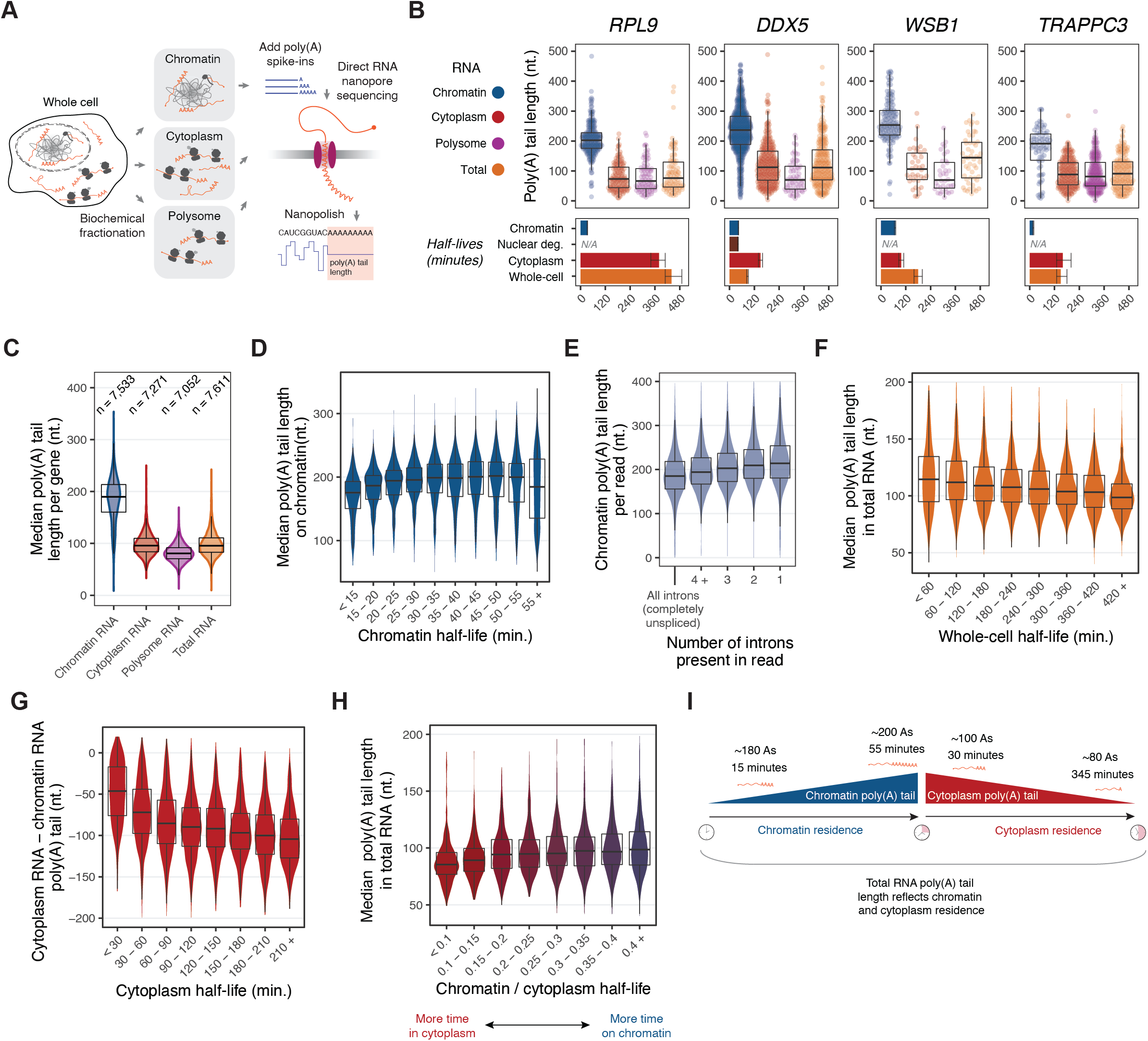
Subcellular compartment-specific poly(A) tail lengths reflect RNA flow rates. (A) Schematic for poly(A) tail length analysis. Chromatin, cytoplasm, polysome, and total RNA from K562 cells were directly sequenced by nanopores. The poly(A) tail length on each RNA was estimated using nanopolish-polya (Workman et al., 2019), and synthetic RNA spike-ins were used to normalize poly(A) tail length across sequencing runs. (B) Poly(A) tail lengths per RNA across compartments for example genes (*RPL9, DDX5, WSB1, TRAPPC3*). Each dot represents an individual RNA. The mean chromatin, nuclear degradation, cytoplasm, and whole-cell half-lives for one replicate are indicated below each gene, with the error bars representing credible intervals. (C) Distribution of median poly(A) tail lengths for each gene covered by >=10 reads in each sample. The number of genes analyzed in each compartment is noted. (D) Median poly(A) tail length for genes covered by >=10 reads in chromatin RNA libraries as a function of chromatin RNA half-life. (E) Poly(A) tail lengths on chromatin as a function of splicing status. Poly(A) tail lengths were analyzed for all chromatin RNA reads with incomplete splicing (containing at least one unexcised intron), binned by the number of introns present. (F) Median total RNA poly(A) tail length for genes covered by >=10 reads in total RNA libraries as a function of whole-cell half-life. (G) The difference between the median chromatin poly(A) tail length and the median cytoplasm poly(A) tail length for all genes covered by >=10 reads in each library, as a function of their cytoplasm half-life. (H) Median poly(A) tail length in total RNA as a function of the relative amount of time spent on chromatin and in the cytoplasm for each transcript, defined as the ratio of chromatin half-life to cytoplasm half-life. Genes covered by >=10 reads in total RNA libraries were included. (I) Model of compartment-specific poly(A) tail lengths with respect to subcellular half-lives. Poly(A) tails of chromatin-associated RNAs lengthen with increased chromatin residence, while cytoplasmic poly(A) tails shorten with increased cytoplasmic residence.

We observed that poly(A) tail lengths gradually shortened as RNAs flowed through the cell, as expected (Figure S7C). These trends could be observed at the single gene level, albeit with variability across genes (Figure 4B, Table S6). For example, median chromatin poly(A) tail lengths ranged from below 200nt to above 250nt (e.g., *TRAPPC3* and *WSB1*, Figure 4B). Globally, the longest poly(A) tails were on chromatin. We measured shorter polyA tail lengths in the cytoplasm, where most genes had median tail lengths of 85–110nt (Figure 4C). Total RNA tail lengths closely resembled those of cytoplasm RNA (Figure 4C). In general, the distribution of tail lengths in polysome RNA were slightly shorter than in cytoplasmic RNA (Figure 4C).

### Poly(A) tail length increases with chromatin residence and decreases with cytoplasm residence

To determine the relationship between poly(A) tail lengths and RNA flow across the cell, we compared the median tail length per gene in each compartment to the respective subcellular half-life. On chromatin, median poly(A) tail lengths were the shortest for genes with chromatin half-lives of less than 15 minutes, and chromatin tail lengths gradually increased with half-life until ~40 minutes (Figure 4D). Across all genes, chromatin poly(A) tails were shortest on completely unspliced transcripts, and tails increased in length on intermediately spliced transcripts as more introns were removed from the pre-mRNA (Figure 4E). This observation indicated that poly(A) tails grow as transcripts spend more time on chromatin, a period also associated with continued splicing. We observed that median cytoplasm RNA poly(A) tail length decreased with longer cytoplasm half-lives in both K562 cells and NIH-3T3 cells (Figure S7E-F), and a negative correlation between median tail length in total RNA and whole-cell residence (Figure 4F), consistent with previous reports (Lima et al., 2017; Subtelny et al., 2014).

Comparing across fractions, we found that transcripts with short cytoplasmic half-lives (<30 minutes) experience a shorting of their polyA tails by ~50 nt, wheras mRNAs with much longer half-lives (>345 minutes) shortened their tails much more by >100 nt (Figure 4G). These observations are consistant with the notion that the tails of transcripts with long half-lives are “pruned” to a certain length, at which point they remain relatively stable (Lima et al., 2017) and findings that transcripts with faster turnover undergo more rapid deadenylation and degradation, without such stabilization at shorter tail lengths (Eisen et al., 2020a).

Based on our findings that chromatin poly(A) tail length increases with chromatin residence and cytoplasm poly(A) tail length decreases with cytoplasm residence, total RNA tail length must be reflective of the relative time spent by transcripts in both compartments. mRNAs that spent relatively more time on chromatin than in the cytoplasm had longer poly(A) tails in total RNA, whereas mRNAs that spent relatively more time in the cytoplasm had shorter tails in total RNA (Figure 4H). For example, *TRAPPC3* and *WSB1* had similar whole-cell half-lives, but *TRAPPC3* spent ~7x more time in the cytoplasm than on chromatin, whereas *WSB1* mRNA spent nearly the same amount of time on chromatin and in the cytoplasm (Figure 4B). Consequently, *TRAPPC3* had a much shorter median poly(A) tail in total RNA (median 89nt) than *WSB1* (median 142nt) (Figure 4B). Thus, median poly(A) tail lengths in total RNA are not only reflective of whole-cell residence times (Figure 4F), but also of the relative times spent on chromatin and in the cytoplasm (Figure 4H-I).

### PUND transcripts are spliceosome targets that exhibit distinct RNA flow, splicing, and poly(A) tail phenotypes

Given the patterns of RBP binding and poly(A) tail lengths across compartments, we wondered whether PUNDs behaved uniquely in any respect. To explore this possibility, we began by comparing the half-lives in each compartment between PUNDs and all other transcripts. In human K562 cells, PUND genes had faster turnover on chromatin, in the nucleus, and at the whole-cell level than genes without evidence of nuclear degradation (Figure 5A, S8A). Notably, PUND genes had longer cytoplasm half-lives than other mRNAs (Figure 5A, S8A), indicating that the transcripts from PUNDs that do get exported are more stable in the cytoplasm compared to transcripts of other genes. Thus, the high nuclear degradation of PUND transcripts is not a trivial reflection of the breakdown of inherently unstable transcripts.

**Figure 5:**
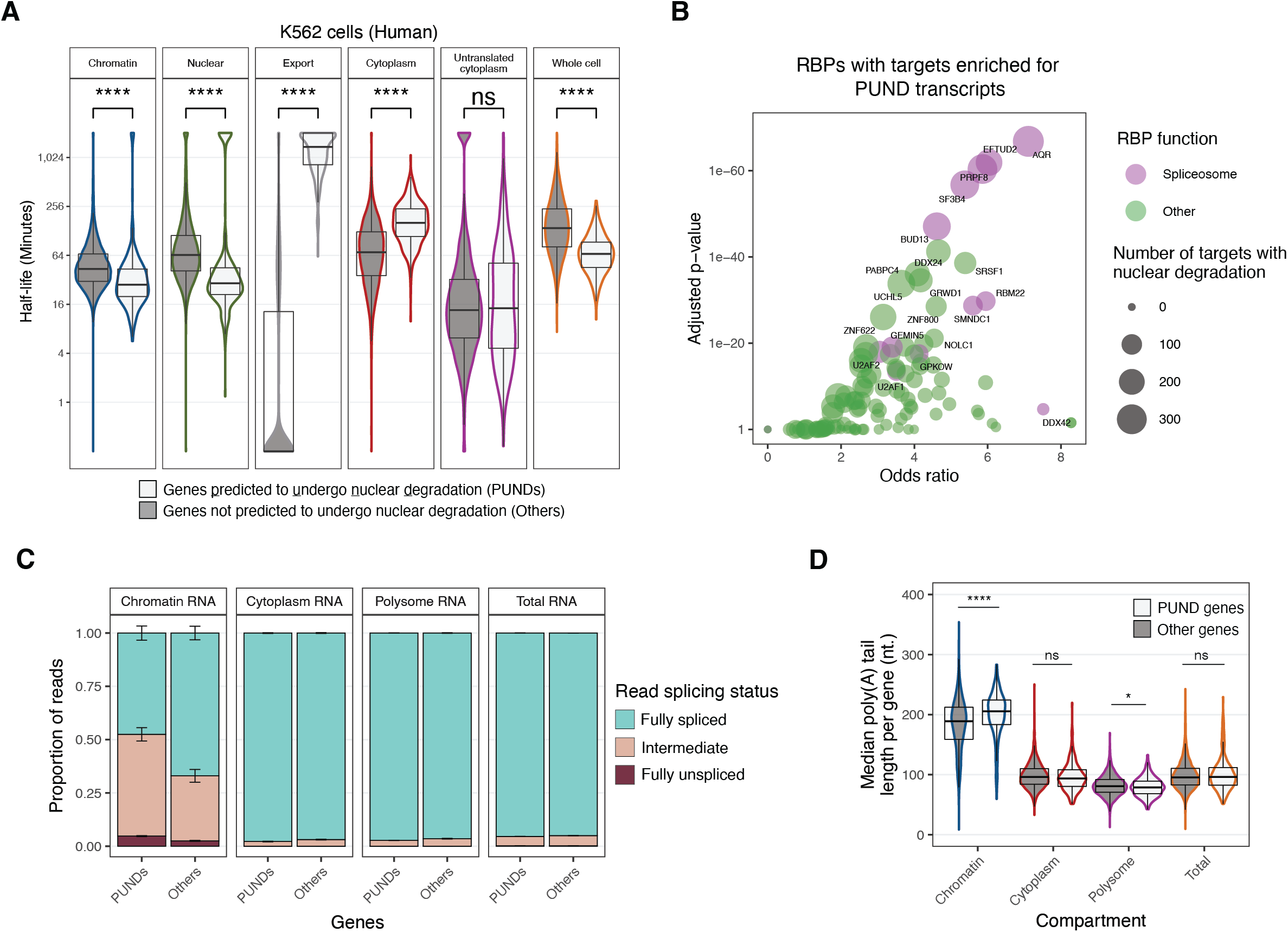
Genes predicted to undergo nuclear degradation (PUNDs) exhibit unique phenotypes related to RNA flow, splicing, and poly(A) tail lengths. (A) Half-lives of all PUND genes (n=408) compared to all other genes in human K562 cells (****: p<0.0001, *: p<0.05, “ns:” not significant, Wilcoxon test). (B) Volcano plot representing the odds ratio and adjusted p-value obtained from a Fisher’s exact test comparing the target and non-target mRNAs for each RBP analyzed in Figure 3 (n=120) to determine which RBPs have enriched PUND targets. RBPs are colored by function as defined by (Van Nostrand et al., 2020), with “other” representing any function that is not “spliceosome.” (C) Splicing levels of PUND and other transcripts in nanopore direct RNA sequencing data across subcellular compartments and in total RNA. Error bars show standard error over two biological replicates. (D) Median poly(A) tail length of PUND genes relative to others across subcellular compartments. The median poly(A) tail length was calculated for each gene covered by >= 10 reads in each sample. Tail lengths were compared between PUND and other genes, significance was noted as in (A).

We found 68 RBPs with target genes significantly enriched for PUNDs (Figure 5B, Table S5). Remarkably, five of the proteins most significantly enriched were components of the spliceosome: AQR, EFTUD2, PRPF8, SF3B4, and BUD13 (Figure 5B). We also detected splicing factor SRSF1 among the most significant RBPs (Figure 5B). In light of this finding, we sought to determine whether PUND transcripts exhibited either slow or fast splicing compared to other mRNAs. Analyses of our direct RNA sequencing libraries revealed PUND genes had more incompletely spliced mRNAs on chromatin than other genes, but overall splicing levels did not differ in other compartments (Figure 5C). Finally, we investigated whether PUND transcripts exhibited any differences in poly(A) tail length relative to other transcripts, and found that they had longer tails on chromatin (median PUND tail length= 210nt, median other= 193nt, p=1.40×10^−16^) (Figure 5D, S8B). This was surprising given that PUND transcripts generally reside on chromatin for less time (Figure 5A), and we observed that transcripts with short chromatin half-lives generally have short poly(A) tails (Figure 4D,I). PUNDs had slightly shorter tails on polysomes than other mRNAs (median PUND tail length=80nt, median other=83nt, p=0.013), but tail lengths did not differ in cytoplasm or total RNA. We conclude that PUND transcripts have more incomplete splicing and longer poly(A) tails on chromatin, but are largely indistinguishable from other transcripts in the cytoplasm, on polysomes, and in total RNA.

### Machine learning model identifies molecular features that explain RNA flow rates

Finally, we sought to identify genetic and molecular features that collectively explain the variability in RNA flow rates (Figure 6A, Table S7). To this end, we developed a LASSO regression model (Figure 6A, S9A) that identifies sparse relevant features through L^1^ regularization (Hastie et al., 2001). The 10× cross-validation and unseen test set performances of our model varied between the subcellular compartments (Figure 6B), ranging from R^2^ = 0.4 for cytoplasmic turnover to R^2^ = 0.13 for polysome loading rates, likely due to the larger uncertainty for these estimates (Figure 1G, S3D). Our model performed as well as the best published whole-cell models when we trained it on our whole-cell rates or on published “ensemble” values (Figure S9B) (Agarwal and Kelley, 2022; Blumberg et al., 2021; Chan et al., 2018; Chen and van Steensel, 2017; Cheng et al., 2017; Sharova et al., 2009; Spies et al., 2013; Yang et al., 2003).

**Figure 6:**
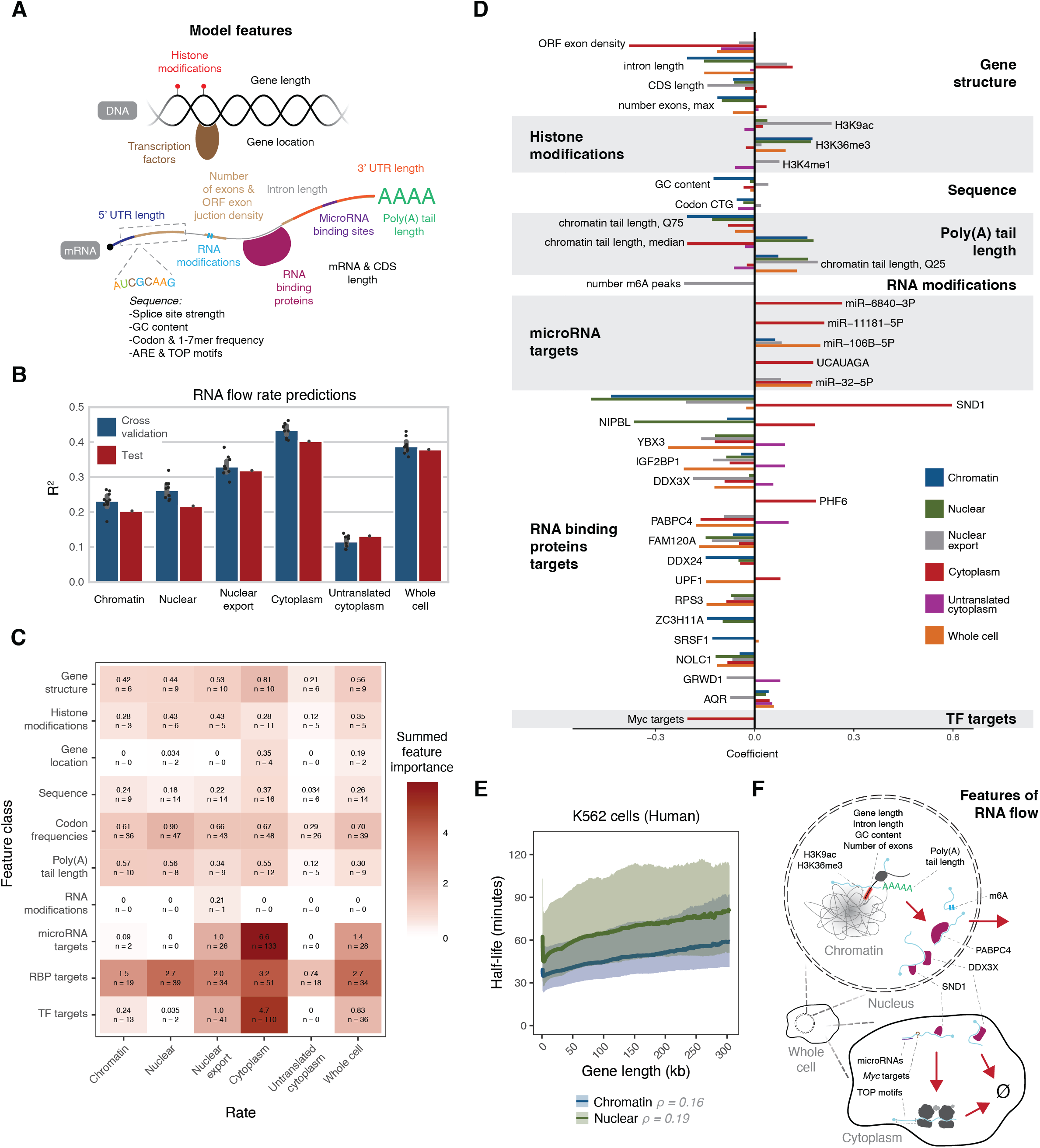
LASSO regression model identifies most relevant genetic and molecular features that predict RNA flow rates. (A) Schematic representing the genetic and molecular features for each gene included in LASSO model (full list: Table S7). (B) 10x cross validation and test set performances of LASSO models predicting subcellular rates. (C) Class feature importances of LASSO models. (D) Top individual features associated with RNA flow. The top 10 features with highest importance of each flow rate were identified and their correlation with any flow rate was shown. Individual features were grouped by feature family as in (C). (E) Continuous averages of chromatin and nuclear half-lives as a function of gene length in human K562 cells. Solid lines represent median half-lives and shaded ribbons represent the third quartile (top) and first quartile (bottom) of half-lives. (F) Schematic depicting relevant features related to RNA flow rates. Features are included in the compartments where they associate with RNA flow rates (top: chromatin, nuclear, and nuclear export half-lives; bottom: cytoplasm and untranslated cytoplasm half-lives).

Across all RNA flow rates, basic gene structure, histone modifications, sequence features, codon frequencies, RBP target sets, and compartment-specific poly(A) tail lengths were the feature classes that provided the most information (Figure 6C, Table S7). We also observed subcellular-specific relevant classes, such as microRNAs promoting cytoplasmic turnover and transcription factors targets associating with nuclear export and cytoplasmic turnover (Figure 6C). Sequence determinants and codon frequencies were collectively relevant as feature classes (Figure 6C), albeit the effect sizes of individual features were generally small and compartment-specific (Figure S9C, Table S7). For example, transcripts containing a 5’ <u>t</u>erminal <u>o</u>ligo-<u>p</u>yrimidine (TOP) motif (Cockman et al., 2020) had faster RNA flow dynamics, including polysome loading rates, but were also more stable in the cytoplasm (Figure S9C, Table S7).

Next, we investigated the strongest individual feature contributions across compartments (Figure 6D). Both intron length and the number of exons were top predictors of chromatin half-lives (Figure 6D). Accordingly, chromatin and nuclear half-lives increased with gene length in both cell lines, likely due to transcription elongation time, albeit with a large variability between genes (Figure 6E, S9E, Spearman correlations for K562: chromatin=0.16, nuclear=0.19, nuclear; NIH-3T3: chromatin=0.20, nuclear=0.15). Gene GC content, histone H3K36 tri-methylation, and SND1 binding all positively influenced chromatin, nuclear, and nuclear export rates (Figure 6D). Consistent with our RBP analysis (Figure 3C-F), DDX3X and PABPC4 were predicted to regulate nuclear export. Notably, the RNA modification N6-methyladenosine (m6A) was also strongly predictive of slower nuclear export rates, but not any other RNA flow rate (Figure 6D).

Our LASSO model predicted that genes targeted by the MYC transcription factor (TF) had long cytoplasm half-lives (Figure 6D), and we confirmed this finding orthogonally by GSEA (Table S4). However, other TFs with fewer target genes, and thus a smaller global contribution, might have remained unidentified in our model. To more sensitively investigate whether more TFs were associated with the RNA flow rates of their target genes, we systematically identified TFs for which the half-lives of targets differed from non-targets (Figure S9F), similar to our RBP target analysis (Figure 3A-B). Remarkably, 193 TFs were associated with altered RNA flow rates (Figure S9F). For instance, targets of the stress-response TFs ATF4 and ATF5 had shorter chromatin half-lives (Figure S9F). Our LASSO model and TF analysis revealed that many uncharacterized Zinc-finger proteins were also associated with altered RNA flow rates (Figure S9F, Table S8). We conclude that many features, underlying a diverse set of regulatory mechanisms, are responsible for subcellular transcriptome dynamics (Figure 6F).

## Discussion

Here we have shown that our analysis of RNA flow, based on subcellular TimeLapse-seq and kinetic modeling, is capable of characterizing the many possible life cycles of mammalian transcripts across the cell and yields subcellular RNA half-lives genome-wide. We observed that the variability in RNA turnover previously observed at whole-cell resolution persisted across all subcellular compartments (Figure 1H). Transcripts with the same whole-cell half-lives flowed throughout the cell at different rates (Figure S4F-G), highlighting the variability in regulation between different genes. In both cell types analyzed, we identified many mRNAs that spent equal or more time on chromatin than in the cytoplasm, even among transcripts with the fastest whole-cell turnover (Figure 2C). We also obtained evidence that many mRNAs remain associated with chromatin for an extended period of time after transcription has completed, during which fully transcribed mRNAs continue to undergo splicing and polyadenylation (Figure 4E).

Subcellular TimeLapse-seq relies on the biochemical purification of subcellular compartments, an operation with inherent limitations. Endoplasmic reticulum (ER) can partially co-sediment with the nuclear fraction in similar cell fractionation methods (Bhatt et al., 2012; Yeom and Damianov, 2017), and if this is occurring in our system, it may explain several minor observations. First, transcripts enriched for functions related to the cell membrane, smooth ER, and COPI-coated vesicles, which are translated at the ER membrane (Fazal et al., 2019; Jan et al., 2014), were enriched in clusters with very long nuclear half-lives (clusters 17-20, n=703 genes, Figure 2B). We also identified a few genes that had extremely long chromatin half-lives but relatively short poly(A) tails (Figure 4D). We hypothesize that the chromatin and nuclear fractions contain a mix of both newly synthesized and preexisting RNAs for these select genes, resulting in artificially long half-lives and likewise, artificially short poly(A) tails. Finally, we are uncertain where various granules and condensates sediment, given their wide range of biochemical properties (Ditlev et al., 2018). Notably, however, P-bodies (monitored by marker protein LSM14A) sedimented in the cytoplasm (Figure 1B) and did not colocalize with polysomes (Figure S1D). In sum, we believe that our biochemical purification accurately assigns the subcellular localizations of most transcripts.

We identified hundreds of genes that are predicted to undergo substantial nuclear degradation in both cell types analyzed where most of the transcripts encoding these genes are never exported from the nucleus. Genes with lower fractions of nuclear degraded transcripts would not be distinguished reliably through our conservative Bayes Factor analysis, so it is likely that more genes exhibit a significant, albeit lesser, degree of nuclear RNA degradation. Indeed, reduction of the Bayes factor criterion from a value of 100 to 10 identifies an additional 98 PUNDs in NIH-3T3 cells and 50 PUNDs in K562 cells. Taking into account the high production rates of many PUNDs, we estimate that at least 20% of all human and mouse protein-coding transcripts are degraded in the nucleus. PUND transcripts have unique features, such as incomplete splicing, long poly(A) tails, and association with splicing factors and other RBPs, that may aid in their identification by nuclear degradation pathways. Given that many PUNDs encode for splicing and nuclear RNA degradation factors, we speculate that nuclear degradation plays a role in nuclear RNA homeostasis (Berry and Pelkmans, 2022; Berry et al., 2022). Overall, we believe nuclear transcript degradation to be pervasive and to likely serve a regulatory function, which is an exciting direction for future work.

We observed significant associations between the rates of RNA flow and many RBPs, and also identified new functions for DDX3X and PABPC4 in regulating nuclear export. This adds to the many known roles of DDX3X, which interacts with multiple nuclear export machineries (Lai et al., 2008; Yedavalli et al., 2004) and forms puncta at nuclear pores (Merz et al., 2007). We report the first nuclear role for PABPC4, which has been shown to shuttle between the nucleus and cytoplasm (Burgess et al., 2011). Collectively, these observations complement the finding that most mRNAs are quickly exported into the cytoplasm following release from chromatin (Figure 1H), but that this step is regulated by DDX3X, PABPC4, and other RBPs.

Our comprehensive analysis of poly(A) tail dynamics across subcellular compartments shows that tail length distributions differ between genes not only in the cytoplasm, but also on chromatin. These results shed new light on the relationship between poly(A) tail length and RNA stability. Although total RNA tail length inversely correlated with whole-cell half-lives in K562 (Figure 4F), this simple relationship does not capture the dynamics across all compartments as cytoplasm half-lives are generally longer than chromatin half-lives (Figure 1H), obscuring the contribution of chromatin RNA poly(A) tails. Indeed, we observed that total RNA tail lengths reflect the ratio of time spent on chromatin and in the cytoplasm for each gene (Figure 4H-I). Thus, we have furthered our understanding of the links between poly(A) tail lengths and mRNA stability to a subcellular resolution, and its full dissection will be enabled by the methodologies introduced here in future studies.

RNA flow represents the cumulative impact of multiple layers of gene regulation that act to control the subcellular fates and trajectories of transcripts. Analysis of RNA flow throughout development and in disease systems is expected to help further elucidate how regulatory programs control cell fate and multicellular phenotypes through subcellular transcript dynamics.

## Acknowledgments

We thank A. Aker, G. Prakash, R.S. Isaac, H. Merens, A. Koenigs, C. Cepko and members of the Churchman lab for helpful discussions; N. Kramer for advice regarding lentiviral transductions; M. Couvillion for creating the SNP-masked genomes; M. Couvillion and A. Sarfatis and for help estimating conversion rates; J. Diego Martin Rufino for providing transcription factor target gene sets; C. Patil, D. Martell, and J. Bridgers for critical reading of the manuscript; the Microscopy Resources on the North Quad (MicRoN) Core at Harvard Medical School for microscopy services; the Bauer Core Facility at Harvard University for sequencing services; the Boston Children’s Hospital Molecular Genetics Core for NanoStrings services. Portions of this research were conducted on the O2 High Performance Computing Cluster, supported by the Research Computing Group, at Harvard Medical School.

## Availability of data and materials

Raw and processed data will be available from GEO. Code for analysis of all data will be made publicly available at http://github.com/churchmanlab/rna_flow (currently private).

## Methods

### Cell Culture

NIH-3T3 cells (ATCC CRL-1658) were maintained at 37°C and 5% CO_2_ in DMEM (ThermoFisher 11995073) with 10% cosmic calf serum (Cytiva SH30087.03), 100 U/mL penicillin, and 100 ug/mL streptomycin (ThermoFisher 15140122). K562 cells (ATCC CCL-243) were maintained at 37°C and 5% CO_2_ in RPMI (ThermoFisher 11875119) with 10% FBS (Corning 35015CV), and 100 U/mL penicillin, and 100 ug/mL streptomycin (ThermoFisher 15140122). HEK-293T cells (ATCC CRL-3216) were maintained at 37°C and 5% CO_2_ in DMEM (ThermoFisher 11995073) with 10% FBS (Corning 35015CV), 100 U/mL penicillin, and 100 ug/mL streptomycin (ThermoFisher 15140122).

### 4sU labeling

Pulse-labeling was performed with 4-thiouridine (4sU, Sigma T4509) resuspended in conditioned cell media. Labeling was performed in NIH-3T3 with cells at 40% confluency (approximately 8×10^6^ cells in a 15cm plate) and a final 4sU concentration of 500uM. Labeling was performed in K562 with 4-5×10^5^ cells/mL at a final 4sU concentration of 50uM. At the beginning of each labeling period, cells were removed from the incubator, 4sU was added directly to the existing cell media, and cells were returned to the incubator for the remainder of the pulse. At the end of the pulse, K562 cells were pelleted at 500xg for 2 minutes. The supernatant (cell media) was discarded for both NIH-3T3 and K562 and cells were immediately placed on ice and fractionated or lysed in 500uL RIPA buffer (ThermoFisher 89900) to collect total RNA.

### SABER-FISH and data analysis

mRNA transcripts corresponding to *Foxo3, Smad3, Gfod1*, and *Myc* were detected in NIH-3T3 cells by smRNA-FISH according to (Kishi et al., 2019). Briefly, 50-80 probes were designed per gene using PaintSHOP (Hershberg et al., 2021) with additional sequences used for SABER appended to the 3’ end in a gene-specific manner (Table S9). All probes corresponding to the same gene were pooled at 10uM in 1x TE buffer (10 mM Tris pH 8.0, 0.1 mM EDTA). Probes were synthesized by first preparing a mix containing 10uL of 5uM hairpin oligo, 10uL of 10x PBS, 10uL of 100mM MgSO_4_, 5uL of dNTP mix (containing 6mM of each A,C,T), 10uL of 1uM Clean.g oligo, 0.5uL of BST enzyme (McLab, BPL-300), and 44.5uL water (see Table S9 for oligo sequences). The reaction mix was incubated at 37°C for 15 minutes and then 10uL of 10uM pooled probes were added. Probes were then concatemerized by incubating at 37°C for 60 minutes before enzyme inactivation at 80°C for 20 minutes. Probes were purified using the MinElute PCR Purification kit (Qiagen 28004) and quantified by Nanodrop (ssDNA setting).

Cells were grown in 8-well poly-L-lysine coated chamber slides (ibidi 80826), labeled with 500uM 4sU for 2 hours, and fixed with 4% PFA in 1x PBS for 10 minutes. Unlabeled cells were also included as controls. Slides were washed 3x 5 minutes in 1x PBST and then incubated in 1x Whyb solution (2x SSC, 1% Tween-20, and 40% deionized formamide) for at least 1 hour at 43°C. 1ug of concatemerized probes were then incubated on slides in 1x Hyb solution (2x SSC 1% Tween-20, 40% deionized formamide, 10% dextran sulfate) for at least 16 hours at 43°C (total volume per ibidi chamber well of 150uL). Slides were wrapped in parafilm and placed in a humidifying chamber within the oven to prevent evaporation. After probe hybridization, each well was washed 2x 30 minutes with 1x Whyb solution (pre-warmed to 43°C), followed by 2x 5 minutes in 2x SSC, 0.1% Tween-20 (pre-warmed to 43°C). Slides were then moved to room temperature and washed 2x 1 minute with 1x PBST (1x PBS, 0.1% Tween). For fluorescent detection of the mRNAs, slides were then transferred to an oven set at 37°C and pre-warmed for 10 minutes. Fluorescent imager oligos were then incubated with the sample at 37°C for 10 minutes, each at 0.2 uM concentration in Imager Hyb (1x PBS, 0.2% Tween). Each well was then washed 3x 5 minutes in 1x PBST at 37°C, before the sample was brought to room temperature.

Samples were then blocked for 1 hour at room temperature in Blocking Solution (1x PBST, 10% Molecular-grade BSA (ThermoFisher AM2616)). Antibodies to detect Lamin B1 and Tubulin were applied in Blocking Solution and incubated for 2 hours at room temperature (see Table S9 for antibody details). Samples were washed 3x 5 minutes with 1x PBST before secondary antibody incubation, which were applied for 1 hour at room temperature in Blocking Solution (see Table S9 for antibody details). Before imaging, samples were washed 3x 5 minutes with 1x PBST and all samples were imaged in 1x PBST.

Images were acquired and puncta were detected according to (West et al., 2022). Briefly, slides were imaged using a Nikon Ti-2 spinning disk inverted microscope with a 40x objective at the Microscopy Resources on the North Quad (MicRoN) core at Harvard Medical School. Images were acquired as multipoint, multichannel images and data were saved and exported as .nd2 files. Images were then split by channel and position and used to generate maximum projections across the z-stacks and a top-hat background subtraction was applied. The nuclear regions within each position were identified by creating a mask from the Lamin B1 signal, and the cytoplasmic regions were similarly identified using Tubulin signal after subtracting the nuclear mask. Finally, mRNA puncta were identified using a Laplacian of Gaussian filter and were called as nuclear or cytoplasmic based on overlap with the masks.

### Cell fractionation

#### Chromatin, nuclear and cytoplasm RNA

Cells were fractionated as per (Danya J Martell, Robert Ietswaart, Brendan M. Smalec, L. Stirling Churchman, 2021). Briefly, cells were lysed in 400uL cytoplasm lysis buffer (10mM Tris-HCl pH 7.0, 150mM NaCl, 0.15% NP-40) and incubated on ice for 5 minutes. Lysate was then layered on top of 500uL sucrose buffer (25% sucrose, 10mM Tris-HCl, 150mM NaCl) and centrifuged at 13,000 RPM at 4°C for 10 minutes to pellet nuclei. The top (cytoplasm) fraction was isolated. Nuclei were resuspended in 800uL nuclei wash buffer (1x PBS with 1mM EDTA, 0.1% Triton-X) and centrifuged at 3,500 RPM at 4°C for 1 minute. To isolate the nuclear fraction, the washed nuclei were resuspended in 500uL RIPA buffer. To isolate the chromatin fraction, the washed nuclei were resuspended in 200uL glycerol buffer (50% glycerol, 20mM Tris-HCl pH 8.0, 75mM NaCl, 0.5mM EDTA, 0.85mM DTT). After resuspension, 200uL nuclear lysis buffer (20mM HEPES pH 7.5, 300mM NaCl, 1M urea, 0.2mM EDTA, 1mM DTT, 1% NP-40) was added and lysates were incubated on ice for 2 minutes before centrifuging at 14,000 RPM at 4°C for 2 minutes. The supernatant was discarded and chromatin pellets were resuspended in 100uL chromatin resuspension solution. The final volume of the chromatin fraction was brought to 250uL with RIPA buffer.

#### Polysome RNA

Cells were lysed in 500uL polysome lysis buffer (25mM HEPES pH 7.5, 5mM MgCl_2_, 0.1M KCl, 2mM DTT, 1% Triton-X, 0.1mg/mL cycloheximide) and incubated on ice for 5 minutes. The lysate was centrifuged at 13,000 RPM at 4°C for 10 minutes to pellet nuclei. The supernatant was then loaded on top of a 12mL 10-50% sucrose gradient (25mM HEPES pH 7.5, 5mM MgCl_2_, 0.1M KCl, 2mM DTT, 0.1mg/mL cycloheximide) and spun in an ultracentrifuge at 35,000 RPM at 4°C for 2 hours. Gradients were fractionated into 13 samples and the RNA absorbance throughout the gradient was monitored with a BioComp 153 gradient station ip (BioComp Instruments, Fredericton, New Brunswick), using a FC-2 Triax flow cell with software v1.53A (BioComp), and fractionated with a Gilson FC203B fraction collector. Lysate from the puromycin-sensitive fractions (Figure S1D) was then pooled as the polysome fraction.

### RNA extraction

RNA extraction was performed using Trizol LS according to the manufacturer’s protocol, except with the addition of DTT at a final concentration of 0.2mM DTT in the isopropanol. For polysome samples, isolated RNA was precipitated using standard ethanol precipitation to reduce the volume of RNA after the Trizol extraction. RNA was quantified using a Nanodrop 2000 (ThermoFisher).

### Western blotting

Samples were mixed at 1:1 volume with 2x Laemmli buffer (4% SDS, 20% glycerol, 0.2M DTT, 0.1M Tris-HCl pH 7.0, 0.02% bromophenol blue), denatured at 95°C for 5 minutes, and kept on ice. Samples were loaded onto a 4-12% Bis-Tris gel (Invitrogen NP0321BOX) in 1x MOPs buffer and run at 160V for 1 hour. The gel was transferred to a nitrocellulose membrane using the wet transfer method in 1x transfer buffer (25mM Tris base, 192mM glycine, 20% methanol) at 400mA for 75 minutes at 4°C. The membrane was blocked in 1x blocking buffer (5% non-fat milk powder in 1x TBST) for at least 60 minutes. Primary antibodies were diluted according to (Table S9) in 1x blocking buffer and incubated with membranes for at least 16 hours at 4°C. Membranes were washed 4x 5 minutes with 1x TBST, incubated for 1 hour at 25°C with secondary antibodies (Table S9), washed again 4x 5 minutes with 1x TBST, and imaged using a Li-Cor Odyssey.

### TimeLapse-seq chemistry and library preparation

Samples were prepared for sequencing according to (Schofield et al., 2018). Briefly, 2.5ug RNA was treated with 0.1M sodium acetate pH 5.2, 4mM EDTA, 5.2% 2,2,2-trifluoroethylamine, and 10mM sodium periodate at 45°C for 1 hour. RNA was then cleaned using an equal volume of RNAClean XP beads (Beckman Coulter A63987) by washing twice with 80% ethanol. The cleaned RNA was then treated with 10mM Tris-HCl pH 7.5, 10mM DTT, 100mM NaCl, and 1mM EDTA at 37°C for 30 minutes. The RNA clean up with an equal volume of RNAClean XP beads was repeated and RNA was quantified by Nanodrop. Library preparation was performed using the SMARTer Stranded Total RNA HI Mammalian kit (Takara 634873) with 0.5-1ug of RNA and samples were sequenced on the NovaSeq (Illumina, San Diego, CA) by the Bauer Core Facility at Harvard University.

### Quantification of newly synthesized RNA from subcellular TimeLapse-seq data

Reads were filtered for quality and adaptor sequences were trimmed using cutadapt v2.5 (Martin, 2011). The first 3nt were trimmed from the 5’ of read1, and the last 3nt were trimmed from the 3’ of read2, corresponding to the 3nts added by the strand-switching oligo during the reverse transcription step in the library preparation. In order to minimize the background mismatch rate, SNP-masked genomes were prepared starting with hg38 and mm10 using non-4sU total RNA TimeLapse-seq reads from K562 and NIH-3T3, respectively (Figure S2B). To prepare the SNP-masked genomes, reads were first mapped to the reference genome with STAR v2.7.3a (Dobin et al., 2013) using parameters --outFilterMultimapNmax 100 --outFilterMismatchNoverLmax 0.09 --outFilterMismatchNmax 15 --outFilterMatchNminOverLread 0.66 --outFilterScoreMinOverLread 0.66 --outFilterMultimapScoreRange 0 --outFilterMismatchNoverReadLmax 1. Variants were then called with BCFtools mpileup (Li, 2011) and call using two bam files as input. The resulting variant call file (VCF) was then split into a file with INDEL records only and a file without INDEL records (substitutions only). The “no INDEL” VCF was further split by frequency of substitution: loci covered by >= 5 reads and with a variant frequency >75% to a single alternate base were assigned the alternate base; loci with variants with an ambiguous alternate base were masked by “N” assignment. The reference FASTA was modified for these non-INDEL substitutions using GATK FastaAlternateReferenceMaker (McKenna et al., 2010). Finally, rf2m (https://github.com/LaboratorioBioinformatica/rf2m) was used with the INDEL-only VCF file to further modify the FASTA genome reference as well as the corresponding GTF annotation file.

The trimmed, filtered reads were aligned to the appropriate SNP-masked genome using STAR v2.7.0a using the following parameters: --outFilterMismatchNmax 15 --outFilterMismatchNoverReadLmax 0.09 --outFilterScoreMinOverLread 0.66 --outFilterMatchNminOverLread 0.66 --alignEndsType Local --readStrand Forward --outSAMattributes NM MD NH. Reads that were not mapped in proper pairs, non-primary and supplementary alignments, and reads aligning to the mitochondrial genome were all discarded using samtools v1.9 (Li et al., 2009). Samples were then grouped by compartment and replicate and converted into .cit files using GRAND-SLAM v2.0.5d (Jürges et al., 2018). Samples were processed through GRAND-SLAM twice, once with the -no4sUpattern option specified and a second time without this parameter. This ensured that the background T>C mismatch rate (p_E_) was calculated using the −4sU sample during the first run and then the data for the −4sU sample was outputted during the second run.

The *a priori* unknown 4sU-induced T>C conversion rate (p_C_) increased with the cellular 4sU concentration throughout the pulse durations (Figure S2A). This resulted in T>C mismatch distributions that differed between genes according to their rates of turnover, such that the assumption of a single global p_C_ for all genes, as in GRAND-SLAM (Jürges et al., 2018), was no longer sufficient. We therefore estimated upper and lower bounds on the gene-specific fractions of new RNA, for each sample as follows. The default GRAND-SLAM output was analyzed to select the 1,000 genes with the fastest turnover, i.e. the 1,000 protein-coding genes with the highest MAP values at the lowest non-0 4sU time point, or the 500 genes with the slowest turnover, i.e. the 500 protein-coding genes with the lowest MAP values > 0.2 at the longest 4sU time point, within each compartment and replicate. Genes with high background T>C mismatches, i.e. MAP in the unlabeled sample >0.05, were excluded from consideration. For each sample, all reads aligning to either the fast or slow turnover set of genes were then analyzed to determine the number of T>C mismatches and total number of T nts across each fragment (considering both read1 and read2 in each pair) using custom scripts. Two dimensional distributions were generated containing the number of fragments with *n* Ts and *k* T>C mismatches, in each of the fast and slow turnover gene group.

To quantify the upper and lower bounds for the 4sU-induced and background T>C conversion rates, we first developed a binomial mixture model for the background conversions with 2 T>C conversion rates (p_E1_ and p_E2_) and a global fraction parameter (π_*E*_) of the two populations, with *Binom* a binomial distribution:

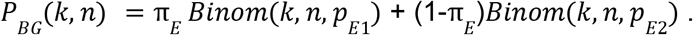

The three parameters in this model were fitted to the above T>C distributions using linear regression (Python v3.7.4, package lmfit v1.0.2, function minimize). We found that our binomial mixture error model better fitted the T>C conversions of the untreated samples (Akaike Information Criterium (AIC): p = 1.0), when compared to the established approach of a binomial error model with a single background T>C conversion (AIC: p < 1e-16) (Jürges et al., 2018). Because p_E2_ turned out to be of similar magnitude (~2%) as the 4sU-induced T>C conversion rate (p_C_, see below), this relatively small second background population ((1-π_*E*_) ~ 2-3%) was essential to include in the model in order to then accurately estimate the 4sU-induced T>C conversion rate (p_C_) and global fraction of newly synthesized RNA (π_*C*_). The 4sU sample T>C distributions were then modeled as the 4sU-induced T>C conversions + background population:

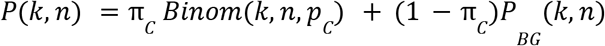

The above fitting procedure was applied to the T>C mismatch distributions of both the fast and slow turnover genes, which generated p_C_HI_ and p_C_LO_, our respective upper and lower bound estimates on p_C_. GRAND-SLAM was run a final two times, once each using p_C_HI_ and p_C_LO_. For each bound *X* ∈ {*HI, LO*}, this resulted in a gene-specific fraction of new RNA (π) posterior distribution: 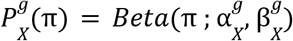, a beta distribution characterized with parameters 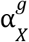 and 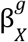. The final gene-specific posterior *P^g^*(*π*) is then as follows:

For the 1,000 genes used to calculate 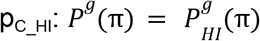.
For the 500 genes used to calculate p_C_LO_ (and those with slower turnover): 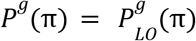.
For the other genes, incorporating the uncertainty over pC, the normalized sum over both bound posteriors: 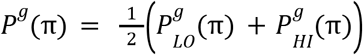

### NanoStrings quantification of new RNA

RNA was denatured at 65°C for 5 minutes and immediately incubated on ice for 2 minutes. The biotinylation reaction was performed with 15ug of denatured RNA in 78ul water, 10ul of MTSEA biotin-XX (Biotium 90066) at 0.25mg/mL in dimethylformamide, and 10uL of 10x buffer (100mM Tris pH 7.5, 10mM EDTA) on a thermoblock at 24°C at 800 RPM for 30 minutes. The biotinylated RNA was purified using Phase Lock tubes (QuantaBio 2302830) using standard chloroform/isoamyl alcohol and ethanol precipitation and quantified by Nanodrop. To synthesize the spike-in RNA, ERCC-00048 DNA with an upstream T7 promoter was cloned into pUC19 and PCR amplified (primers in Table S9) with Phusion polymerase (New England Biolabs M0530S) using the following cycling conditions: 98°C for 30 seconds, then 35 cycles of 98°C for 10 seconds, 61°C for 15 seconds, and 72°C for 30 seconds, followed by a final extension of 72°C for 2 minutes. The PCR product was cleaned using the Monarch® PCR clean up kit (New England Biolabs T1030S) according to the manufacturer’s protocol, quantified by Nanodrop, and used as the template for *in vitro* transcription reaction (New England Biolabs E2040S) performed according to the manufacturer’s protocol. RNA was purified using standard ethanol precipitation and quantified by Nanodrop.

3ug of biotinylated RNA and 60pg of *in vitro* transcribed spike-in RNA (ERCC-00048) in 200ul water was mixed with 100ul of beads from the uMACS Streptavidin kit (Miltenyi Biotec 130-074-101) on a thermoblock at 24°C at 800 RPM for 15 minutes. The columns were washed once with 900uL of wash buffer (100mM Tris pH 7.5, 10mM EDTA, 1M NaCl, 0.1% Tween-20). The RNA/bead mixture was then passed through the washed columns twice and the flow-through RNA was collected, purified using the miRNeasy Nano kit (Qiagen 217084) according to (Schwalb et al., 2016), and quantified by Nanodrop. 150ng of RNA was hybridized with gene-specific DNA probes (Table S9), XT Tagset-24 capture and target probes (NanoString Technologies, Seattle, WA), and hybridization buffer (NanoString Technologies, Seattle, WA) according to the manufacturer’s protocol at 67°C for at least 16 hours before being loaded onto a nCounter Sprint Cartridge and quantified using the nCounter SPRINT Profiler (NanoString Technologies, Seattle, WA) at the Boston Children’s Hospital Molecular Genetics Core. The fraction of new RNA was calculated at each time point according to the following equation:

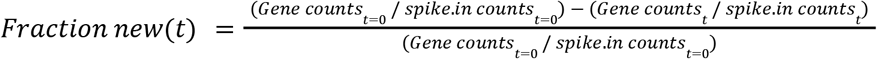

Any negative fraction of new RNA values were replaced with 0s. Note that *JUN* and *SHOX2* were included in the probe set but not analyzed due to low RNA counts (within the range of the included manufacturer’s negative controls) across most time points. Non-coding transcripts *MALAT1* and *COX1* were also included in the probe set but not analyzed for these experiments.

### Kinetic modeling of RNA flow

The kinetic model (Figure 1E), defined by a system of coupled ordinary differential equations (ODEs, Figure S3A), describes the average time evolution of the variables, i.e. the 4sU-labeled RNA levels in their respective subcellular compartments at 4sU-pulse time *t* : T, whole-cell (<u>t</u>otal); CH, <u>ch</u>romatin; N, <u>n</u>ucleus; CY, <u>cy</u>toplasm; P, <u>p</u>olysome; M, (<u>m</u>ature) untranslated cytoplasm. In addition to the RNA flow rates (vector 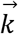, Figure S3A), the ODEs also contain the *a priori* unknown RNA production rate *k_P_*. All rates are considered unaffected by 4sU treatment, a model assumption for which we provide evidence (Figure S1B-C). To solve these ODEs analytically (Adams and Essex, 2021), we set all labeled RNA levels to zero at t=0, i.e. before any 4sU pulsing (boundary conditions). Next, the integrating factor method was used to obtain solutions for the fractions that only depend on a single RNA flow rate: T(*k_WC_, k_P_, t*), CH(*k_CH_, k_P_, t*) and N(*k_N_*, *k_P_, t*). Inserting these expressions then enabled solving the coupled ODEs of 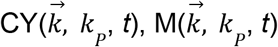 and 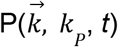, again using an integrating factor. Since the observed quantities are fraction of new RNA, rather than RNA levels we derived the model fraction of new RNA for each compartment 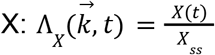, with *X_SS_* levels of all (labeled and unlabeled) compartment RNA, which equals the steady state solution to the ODE, i.e. when 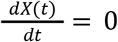. Both *X_SS_* and *X*(*t*) are linear in *k_P_*, so 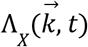 no longer depends on *k_P_*. Full expressions of the 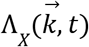 solutions for all compartments are available at https://github.com/churchmanlab/rna_flow/new_total_ratio_jit.py.

As described in section *Quantification of newly synthesized RNA from subcellular TimeLapse-seq data* (Figure S2A), the gene-specific (g) fraction of new RNA (π) Posterior probability density function, 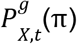, was experimentally estimated for 4 different 4sU-pulse times (t = 15, 30, 60 and 120 minutes) for each compartment *X*, through GRAND-SLAM’s Bayesian inference framework (Jürges et al., 2018). To analytically derive the (in some cases multivariate) posterior on the RNA flow rates (Adams and Essex, 2021), we equated our model 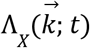 to π, multiplied the posteriors from all (independent) timepoints, and applied the calculus of multivariate change of variables from π to 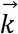 (see resulting expression in Figure S3A).

For the cases with a univariate RNA flow rate posterior (compartment X = T, CH, or N), this immediately provided the posterior distribution on 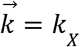. Summary statistics (MAP, Mean and 95% CIs) of the posterior were then determined as follows. Through numerical evaluation of the posterior on a grid (N=1,000 data points) over the prior rate domain [10^−4^, 10^4^] (unit: min^−1^), the MAP (Python package numpy v1.16.5, function: argmax) and 95% CIs were determined (see expressions in Figure S3A). The mean rate (expression in Figure S3A) was obtained through numerical integration (Python package: scipy v1.6.2, function integrate.quad and dblquad). To speed up the numerical integration calculations, integrand functions were coded with “just in time” compilation (Python packages: numba v0.53.1, numba-scipy v0.3.0, function jit). The mean is preferred when using a single number as the RNA flow rate estimate, because it considers all of the posterior distribution. Summary statistics in the results section were reported as the mean between both biological replicates unless noted otherwise. The MAP and 95% CI together provide a more fine-grained characterization.

For the multivariate posteriors (compartment: CY, P), we marginalized the posterior by integrating over the 95% CIs of the already determined upstream RNA flow rate(s) (Figure S3A). For example P_CY_ depends on 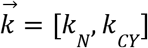, and the *k_N_* 95% CI was already obtained through the univariate procedure described above, so *k_N_* was integrated out, resulting in the posterior on *k_CY_*. Then, posterior summary statistics were calculated as described above.

Lastly, the nuclear export rate *k_E_* was determined (Figure S3A). Deterministically, the time duration of nuclear export equals the difference between the nuclear and chromatin residence times: 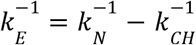. Using its probabilistic analog (Adams and Essex, 2021), the export posterior is then obtained by numerical integration (as described above) over the convolution of the nuclear and chromatin posteriors (see resulting expression in Figure S3A).

Deterministically, the whole-cell RNA levels equal the sum of nuclear and cytoplasmic RNA: T(t) = N(t) + CY(t) (Figure S3A). In absence of nuclear RNA degradation, the whole-cell half-life thus equals the sum of nuclear and cytoplasmic half-lives (Figure S4H-I). In presence of nuclear degradation (Table S2, see the next section for the PUND identification procedure), this simple relation between half-lives no longer holds (Figure S4H-I).

Furthermore, for PUNDs the above nuclear export rate posterior is not appropriate, because that calculation assumes no nuclear degradation is occurring. Deterministically, when including a nuclear degradation rate in the model (Figure 1E), the nuclear turnover rate becomes the sum of nuclear degradation and export rates (Figure S3A): 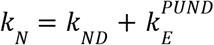. This minimal model extension does not consider the chromatin compartment explicitly. However, in the limit that nuclear export after release from chromatin is much faster than the other rates, the above expression becomes an exact equation where 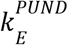 represents the rate of release from chromatin (plus nuclear export) if not nuclear degraded, whilst *k_ND_* equals the nuclear degradation rate, which can occur on chromatin and in the nucleoplasm. Indeed, this limit assumption is consistent with our observations, because for PUNDs the observed chromatin and nuclear turnover rate distributions are similar (Figure 5A). To estimate 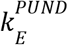, we first derived the deterministic nuclear degradation model whole-cell new RNA levels 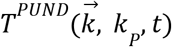 in terms of 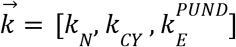 (Figure S3A), and used our approach above to now derive the whole-cell multivariate posterior over 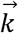, followed by marginalization over *k_N_* and *k_CY_*, to obtain the 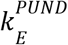 posterior. Summary statistics were then determined as described above. Lastly, given the nuclear posteriors over *k_N_* and 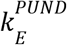, the posterior over 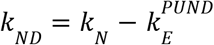 was then derived (see expression in Figure S3A) and calculated through numerical integration (as described above).

Least squares estimation (LSE) of the RNA flow rates acted as a simple deterministic comparison model for the above described Bayesian probabilistic model. LSE was performed by fitting the model 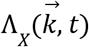 to the MAP(π) timecourse values (scipy.optimize.least_squares, arguments: bounds=[10^−6^,∞], gtol=1e-14, ftol=1e-14, loss=‘linear’). For the multivariate cases, the stepwise estimation approach was used again, as described above. For example, for the cytoplasm, the LSE nuclear turnover rate estimate 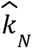 was inserted to enable fitting of 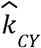. The LSE model suffers from the limitation that the uncertainty on the π estimates and upstream RNA flow rates is not taken into account. For the polysome compartment, this meant that no reproducible *k_PL_* estimates could be obtained with the LSE model, in contrast to our Bayesian model (Figure S3E). Because of the same drawback, the LSE, but not the Baysian model, also predicted a number of biologically unlikely fast rate values (Figure S3E). Besides these differences, we generally observed a strong correspondence between our Bayesian MAP and LSE rate estimates (Figure S3E). In addition to these “best fit” estimates, provided by both models, only the Bayesian model provides a full posterior distribution over the rate domain (Figure 1G), and thus also a 95% CI (Figure S3E), which indicates the range of rate values consistent with the subcellular Timelapse-seq data.

Since the NanoString approach provides single fraction of new RNA values, as opposed to a posterior distribution, NanoStrings RNA flow rate estimation was performed with the LSE model, as described above.

### Bayes Factor model comparison to identify nuclear RNA degradation

To perform formal Bayesian model comparison, we calculated the Bayes factor *K* (Kass and Raftery, 1995), i.e. the ratio of likelihoods of the kinetic model that includes a nuclear degradation rate (alternative hypothesis *M_1_*) over the simpler “nuclear residence” model with no nuclear degradation (null hypothesis *M_0_*): 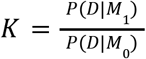. *D* indicates the data, in this case subcellular Timelapse-seq used to distinguish the two models: the timeseries of nuclear, cytoplasmic and whole-cell fraction of new RNA posteriors (Figure S2B), as described in the above sections. The likelihood for the nuclear residence model for gene g is then:

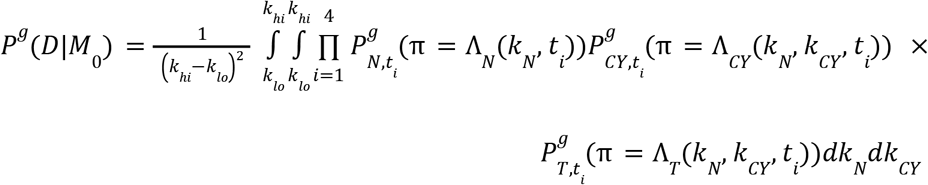

And the equivalent for the nuclear degradation model:

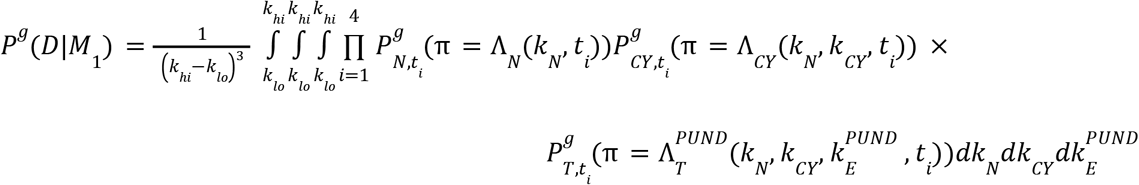

*k_lo_*=10^−4^ and *k_hi_*=10^4^ min^−1^ indicate the prior rate domain bounds. Note that, although the nuclear degradation rate is not explicitly present in the above equation, it is still included in this model since by definition 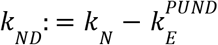. We calculated these integrals numerically (as described in the previous section).

When *K* > 100, it is considered “decisive” evidence in favor of the alternative model, and “strong” evidence if *K* ranges from 10 to 100 (Kass and Raftery, 1995). In our case, genes with transcripts <u>p</u>redicted to <u>u</u>ndergo <u>n</u>uclear RNA <u>d</u>egradation (PUNDs) are defined as having *K* > 100 for both biological replicates (Table S2). Bayes factors and model likelihoods for all genes and replicates are included in Table S1.

Lastly, for each PUND gene, we estimated *f_ND_*, the average fraction of transcripts that are nuclear degraded as opposed to exported: 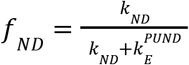, using the mean nuclear degradation (*k_ND_*) and export 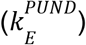 rates as described in the previous section. With these fractions, we then estimated the total cellular fraction of nuclear degraded protein-coding transcripts as: 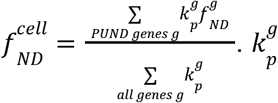 indicates the gene-specific steady state mRNA production rate (unit: RPKM min^−1^), as estimated from the chromatin compartment RNA levels (units: RPKM) and our chromatin turnover rates: 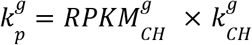. We also confirmed our conclusions were consistent when using RNA production rate estimates derived from our whole-cell Timelapse-seq data. Notably, whole-cell estimates suffer from the drawback that whole-cell turnover rates are a convolution of nuclear and cytoplasmic degradation rates in the case of PUNDs, which then underestimate the resulting production rate estimates. Using the chromatin compartment data resolves this bias.

### Hierarchical clustering of genes by RNA flow rates

Hierarchical gene clustering (scipy.hierarchy.linkage, arguments: metric =‘seuclidean’, method=‘complete’, optimal_ordering=False) was performed on log-transformed half-lives, i.e. 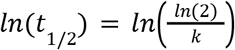, with *k* the MAP and 95% credible interval (CI) endpoints of the turnover rate posteriors, for two biological replicates and all the subcellular compartments: chromatin, nucleus, cytoplasm, polysome and whole-cell. Genes with missing half-live values were excluded. Other RNA flow rates, i.e. nuclear export and degradation, are derived from these subcellular compartment posteriors, and therefore not used for clustering, but nevertheless included in the heatmap for visualization (Figure 2B). For the polysome compartment, only the 95% CIs were used, given that their MAP values were less reproducible across biological replicates (Figure S3D). To identify gene clusters (scipy.hierarchy.fcluster, arguments: criterion=‘maxclust’), we allowed sufficient granularity through a total cluster number of 50. To ensure the robustness of our findings (Figure 2B), the following downstream analyses were also repeated with a total cluster number of 35 and 100. Next, we filtered out all clusters that comprised of genes with irreproducible patterns, i.e. if the median of the half-lives within a cluster differed at least 4 fold between biological replicates for any of the compartments. This resulted in 27 clusters with reproducible patterns, comprising 10,326 genes. Lastly, we reordered the genes from short to long half-lives, whilst respecting the hierarchical clustering (R version 4.1.1, package pheatmap v1.0.12, function: reorder, arguments: agglo.FUN = mean), after which the heatmap with clustered half-lives was visualized (package pheatmap, function: pheatmap, Figure 2B).

### Functional analysis of genes clustered by RNA flow rates

Gene Ontology enrichment analysis of each human gene cluster was performed in Python (package: GOAtools v1.1.6, object: GOEnrichmentStudyNS, arguments: propagate_counts = True, alpha = 0.05/27, to correct for the multiple testing over all 27 clusters, methods = [‘fdr_bh’], Benjamini Hochberg multiple testing correction over the GO terms, and gene_universe = all human genes with any RNA flow rates) (Klopfenstein et al., 2018). For each cluster, the most enriched GO term, i.e. with highest odds ratio (Figure 2B), and all significant GO terms are listed (Table S3). GO enrichment analysis of the human and mouse PUND genes was performed as for the gene clusters, except with alpha=0.05, and for mouse PUNDs with mouse GO annotations and mouse-specific gene_universe (Figure 2A, S5A, Table S3).

Gene set enrichment analysis (GSEA) (Subramanian et al., 2005) was performed with MSigDB (v7.5.1) gene sets h.all, c5.all, c3.all and c2.all (Liberzon et al., 2015), and parameter settings as in (Ietswaart et al., 2021). Briefly, GSEAPreranked (v4.2.2) was run with ranked, Z-score normalized, log-transformed mean half-lives of each human RNA flow rate type as .rnk input file (Table S4).

### RNA binding protein associations with RNA flow rates

K562 eCLIP data for a total of 120 RBPs from (Van Nostrand et al., 2020) was analyzed. The following RNA binding proteins were excluded from the following analyses due to extremely low number of target genes: SLBP, SBDS, UTP3, SUPV3L1, WDR3, PUS1, GNL3, and RPS11. To identify RBPs with significant associations with RNA flow, each replicate of the K562 RNA flow rates were analyzed independently. For each RBP and subcellular half-life, genes were identified as “targets” if the gene contained at least one eCLIP peak with significant enrichment over input. The half-lives of target genes were compared to the half-lives of “non-target” genes (those lacking any significant eCLIP peaks) using a Wilcoxon test with Bonferroni multiple testing correction. The target/non-target half-life was quantified by dividing the median target half-life / median non-target half-life. To identify RBPs containing targets significantly enriched for PUND genes, the targets and non-targets were defined as above. Enrichment was quantified by performing a Fisher’s exact test with Bonferroni multiple testing correction.

### shRNA knockdown of DDX3X and PABPC4

K562 knockdown lines were generated according to (Sundararaman et al., 2016) with slight modifications. Plasmid DNA was purified from pLKO.1 backbone vectors expressing shRNAs targeting DDX3X (Horizon Discovery, TRCN0000000003), PABPC4 (Horizon Discovery, TRCN0000074658), and a scrambled control (Addgene 1864). In parallel, psPAX2 (Addgene 12260) and pMD2.G (Addgene 12259) lentiviral plasmid DNA was purified. All plasmid DNA was quantified by Nanodrop. HEK-293T cells growing in 6-well plates at 50% confluency were transfected with 500ng of shRNA-expressing plasmid, 500ng of psPAX2 plasmid, 50ng of pMD2.G plasmid, and 3.1ul FuGENE HD transfection reagent (Promega E2311) in a total volume of 100ul with Opti-MEM I media (ThermoFisher 31985062). Media was discarded after 24 hours and lentiviral-containing media was collected at 48 and 72 hours after transfection (replacing media every 24 hours) and stored at −80°C. Lentiviral transduction was performed by combining 2×10^6^ K562 cells in 1.75mL media, 1.25mL thawed lentiviral-containing media, and 24ug polybrene (Sigma TR-1003-G). Cells were centrifuged at 1,000 RCF at 33°C for 2 hours, the supernatant was discarded, and replaced with 3mL of K562 media. After 24 hours, 3ug/mL puromycin (Sigma P9620) was added and cells were maintained in the presence of the antibiotic at 0.2-1.0×10^6^ cells/mL for 4 days. Knockdowns were confirmed by western blotting analyses.

### Poly(A) selection and direct RNA sequencing

Poly(A)+ selected from 15-30ug of RNA using the Dynabeads mRNA purification kit (ThermoFisher 61006) according to the manufacturer’s protocol and quantified by Nanodrop. Synthesis of yeast spike-in RNAs was modeled after the protocol described in https://www.ebi.ac.uk/ena/browser/view/PRJEB28423?show=reads for the *S. cerevisiae ENO2* gene. Briefly, six different *S. cerevisiae* genes (*BDC1, ICT1, HIF1, ENO2, YKE4, HMS2*) were amplified from their genomic locus using HiFi Hotstart DNA polymerase (KAPA) (primers in Table S9) in a total volume of 100 uL using the following cycling conditions: 3 minutes at 95°C, then 30 cycles of 15 seconds at 95°C, 15 seconds at 62°C, 2 minutes at 72°C. The PCR amplicons were purified using 1X volume RNA Clean XP beads and eluted in 33 uL water. A second round of PCR was performed with nested primers, wherein the forward primer encodes a T7 RNA polymerase promoter site and the reverse primers have either 10, 15, 30, 60, 80, or 100 thymidines on the 5’ end (primers in Table S9) using the following cycling conditions: 3 minutes at 95°C, then 18 cycles of 15 seconds at 95°C, 15 seconds at 62°C, 2 minutes at 72°C. The PCR amplicons were purified using 1X volume RNA Clean XP beads and eluted in 33 uL water. In vitro transcription was performed using 500ng of DNA template and the MEGAScript™ T7 Transcription kit (ThermoFisher AM1334) according to the manufacturer’s instructions. RNA was cleaned up with the MEGAClear™ Transcription Clean-up kit (ThermoFisher AM1908) according to the manufacturer’s instructions, the concentration was measured by Nanodrop, and the size of the transcripts was verified by TapeStation (Agilent). The six transcripts were pooled at an equimolar concentration (10 picomoles each). 400-700ng of poly(A)+ RNA was combined with 5% spike-in RNA and used to generate direct RNA sequencing libraries with the SQK-RNA002 kit (Oxford Nanopore Technologies) according to the manufacturer’s protocol, except for the ligation of the reverse transcription adapter (RTA), which was incubated for 15 minutes instead of 10 minutes. Samples were sequenced on a MinION device (Oxford Nanopore Technologies) with FLO-MIN106D flow cells for up to 72 hours with live basecalling using MinKNOW.

### Direct RNA sequencing data analysis

All reads with a base calling threshold >7 were converted into DNA sequences by substituting U to T bases. Reads were aligned to the reference human genome (ENSEMBL GRCh38, release-86) concatenated with the six yeast spike-in sequences using minimap2 (version 2.10-r764-dirty) (Li, 2011) with parameters -ax splice -uf -k14. Poly(A) tail lengths were estimated using nanopolish v0.13.3 (Workman et al., 2019). Raw signal fast5 files were indexed with nanopolish index and poly(A) tail lengths were calculated with nanopolish polya using default parameters. Reads with the quality control flag “PASS” and with estimated tail lengths greater than 0 were used in subsequent analyses. To map aligned reads to annotated genes from ENSEMBL GRCh38 (release-86), we used bedtools intersect (Quinlan and Hall, 2010) with options -s -F 0.5 -wo -a $ensembl_bed_file -b $bam, requiring that at least half of the read map to a given gene. For subsequent poly(A) tail length analyses, we filtered for protein-coding genes with at least 10 mapped reads in each sample.

For normalization of poly(A) tail lengths to the spike-ins, we used a median of ratios strategy modeled after the size factor calculation for differential gene expression in DESeq (Love et al., 2014). Poly(A) tail lengths from reads mapping to the yeast spike-in sequences were extracted. For each spike-in, the median poly(A) tail length was calculated in each sample and the geometric mean of medians across samples was computed. The ratio of the median poly(A) tail length per sample over the geometric mean was calculated. Finally, the size factor was defined as the median of ratios across the six spike-ins in each sample. Poly(A) tail lengths from endogenous genes were divided by this size factor for each read, yielding the normalized poly(A) tail length. Of note, the size factors ranged between 0.95 and 1.02 (Figure S7B), indicating low technical variability between sequencing runs.

Analysis of RNA 3’ ends (deriving from the 50 ends of sequenced reads) was performed as described in (Drexler et al., 2020, 2021). Briefly, “Poly(A)” sites are defined as regions within 50 nucleotides of the end coordinate of annotated protein-coding genes or RNA-PET annotations from cytoplasm and chromatin fractions in K562 ENCODE data (ENCODE Project Consortium, 2012). Determining the splicing status of introns and reads was performed as described in (Drexler et al., 2020, 2021). Code for the analysis of RNA 3’ ends and determining the splicing status of introns and reads are available at https://github.com/churchmanlab/nano-COP.

### LASSO machine learning model for RNA flow rate determinants

The objective was to develop a machine learning model that explains a gene-specific RNA flow rate value in terms of that gene’s molecular and genetic features (Figure S9A). Given the large number of input features (70145, Table S7), LASSO regression was chosen as a model because it is a linear model, *y* = β*X*, with L1 regularization, which ensures sparse feature selection (Hastie et al., 2001): 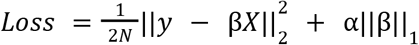 (Python, package scikit-learn v1.0.1, function linear_model.Lasso, arguments: fit_intercept=False, random_state=42, selection=‘random’, max_iter and alpha as specified below). RNA flow rates with genome-wide coverage, i.e. chromatin, nuclear, cytoplasmic, untranslated cytoplasm, and whole-cell turnover and nuclear export, were log-transformed, followed by Z-score normalization, resulting in the model dependent variable (*y*). Rates from different genes constitute independent data points used for model training (*N*). The input features originated from various sources and were grouped into classes based on their biological type (Table S7). Features classes gene location, histone modifications, microRNA targets, RBP targets, and TF targets correspond to gene sets, i.e. categorical features that were one hot encoded into the LASSO feature matrix (*X*). All other features were quantitative, and therefore Z-score normalized to facilitate regularization.

To learn the relevant features, and their effect sizes (coefficient vector β), that best explain the rate variation across the genome, we took a two step learning approach. We split the rates into a 90% training set and a 10% test set, with an identical split between biological replicates. Next, we performed the following round 1 LASSO for each feature class separately (Figure S9A): (1) Using the features from an individual class, we performed 10x cross validation (CV) twice, once on the training rates from each biological replicate, for a range of values of hyperparameter α : [10^−4^, 10^−3^, 10^−2^, 10^−1^] if the number of features < 1000, or else: [10^−3^, 10^−2^, 10^−1^], and max_iter_ = 2e4. (2) The optimal (round 1) α was then identified, such that the 10x CV R^2^ distribution, joined over both replicate runs, was significantly larger than zero in a one-way t-test (p < 0.05, scipy.stats.ttest_1samp, arguments: popmean=0, nan_policy=‘omit’, alternative=‘greater’) and larger than any previously selected α in a two-way t-test (p < 0.05, scipy.stats.ttest_rel, arguments: nan_policy=‘omit’, alternative=‘greater’), similar to a selection approach by (Agarwal and Kelley, 2022). If the average performance did not exceed 0 for any α, none of the features were selected for LASSO round 2 from that particular class. (3) Given the optimal α values for each replicate, any individual feature i with a model coefficient β_*i*_ > 0 in both replicate runs, were thus reproducible and (round 1) relevant and thus selected for round 2 learning.

Next, round 2 LASSO was performed: (1) Reproducible round 1 relevant features from all classes were merged into one feature matrix. (2) The training set rates from both biological replicate were joined into the same 10x CV data split to increase the amount of training data. (3) 10x CV was then performed with a fine-grained range for α: [10^−4^, 3.3×10^−4^, 6.6×10^−4^, 10^−3^, 3.3×10^−3^, 6.6×10^−3^, 10^−2^, 3.3×10^−2^, 6.6×10^−2^, 10^−1^] and max_iter_ = 5e4. (4) The final optimal α was then identified by finding the maximal average 10x CV R^2^, such that the average 10x CV prediction R^2^ did not exceed the average 10x CV training R^2^ by more than 10% (Figure 6B, S9B), to avoid overfitting and robust identification of relevant features. (5) Given the optimal α value, any feature i with a (round 2) model coefficient β > 0 was considered a relevant feature (Figure 6C-D, S9C-D, Table S7). (6) Lastly, we tested the trained round 2 LASSO model by determining its performance on the 10% unseen test set (Figure 6B, S9B).

For the “consensus” whole-cell half-lives (Agarwal and Kelley, 2022), the 10x CV and testing was performed as in round 2 LASSO, described above, with reproducible round 1 relevant features from our whole-cell turnover rates (Figure S9B).

Continuous averaging plots (Figure 6E, S9E), were generated as in (Ietswaart et al., 2017), with minor modifications. First, genes were ranked *g* = 1.. *N* according to their gene length from short to long, where *N* is the total number of genes. This was followed by calculation of the “continuous” averages, < *L_k_* > with *k* = 1.. (2*N* – 1), over these ranked gene subpopulations of their gene length *L*, i.e. the independent variable: 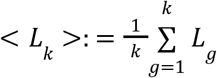 and 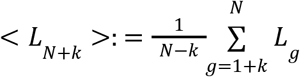. Next, the corresponding continuous median, and Q25 and Q75 (shaded error bands) of the hal-lives, i.e. dependent variables, were calculated over the same gene subpopulations. The shortest 1% and longest 1% of genes were excluded from this analysis.

### Transcription factor associations with RNA flow rates

The same transcription factor target gene sets were used as the input features in our LASSO model (Table S7). The statistical analysis was performed as for the RBP targets as described above in *RNA binding protein associations with RNA flow rates*.

## Supplementary Tables

**Table S1:** RNA flow rates of protein-coding genes in human K562 and mouse NIH-3T3 cells.

**Table S2:** Gene lists generated and used in this study.

**Table S3:** Gene ontology (GO) enrichment results in human K562 and mouse NIH-3T3 cells.

**Table S4:** Gene set enrichment analyses (GSEA) of each RNA flow rate with genes ranked by subcellular half-lives in human K562 cells.

**Table S5:** RNA binding proteins significantly associated with RNA flow rates in human K562 cells.

**Table S6:** Normalized median subcellular and whole-cell poly(A) tail lengths of protein-coding genes in human K562 cells.

**Table S7:** LASSO model features and performances.

**Table S8:** Transcription factors significantly associated with RNA flow rates in human K562 cells.

**Table S9:** Primers, antibodies, and plasmids used in this study.

---

**Supplemental Figure 1:**
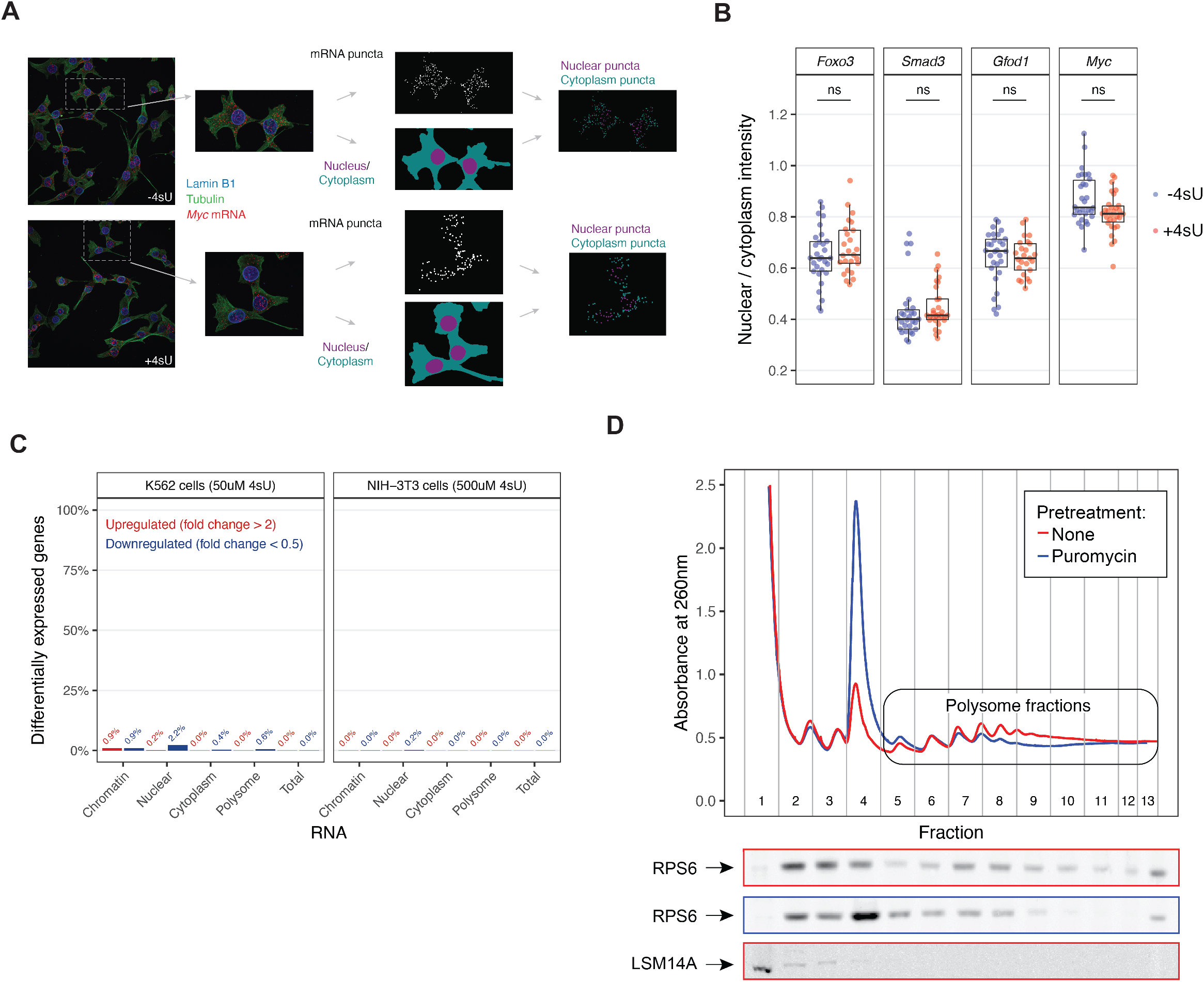
Optimization of biochemical fractionation and 4-thiouridine labeling conditions used in subcellular TimeLapse-seq, related to Figure 1. (A) SABER-FISH was performed in mouse NIH-3T3 cells to visualize *Myc* mRNA according to (Kishi et al., 2019) following 2 hours of 500uM 4sU treatment relative to no 4sU treatment. In addition to hybridizing *Myc* mRNA probes, concurrent immunohistological staining of Lamin B1 and alpha-Tubulin was performed to stain the nucleus and cytoplasm, respectively. mRNA puncta were identified according to (West et al., 2022), and nuclear and cytoplasmic regions within the image were segmented (see Methods for details). Using these defined regions, mRNA puncta were identified as nuclear or cytoplasmic and the puncta intensity within each compartment was summed over all cells in each field of view. A total of 25-30 fields of view were analyzed for all genes. (B) Summary of data for all genes (*Myc, Foxo3, Smad3, Gfod1*) analyzed according to (A). A t-test was performed to compare the differences in intensities between 4sU-treated and control cells (“ns:” non-significant). (C) Number of differentially expressed genes across subcellular compartments in cells following 2 hours of 4sU pulse-labeling relative to no labeling. Differentially expressed genes were defined as those with fold change>2 or <0.5 with an adjusted p-value of <0.01 when comparing RNA-seq read counts to the unlabeled samples for each compartment using DESeq2 (Love et al., 2014). (D) Purification of actively translating ribosomes by sucrose density gradient ultracentrifugation. Polysome profiling traces of K562 cell lysate were measured following no drug treatment (red) or 1 hour of 100ug/ml puromycin treatment (blue). Each fraction was also analyzed by western blotting for a ribosomal protein (RPS6) and a P-body component (LSM14A). Fractions 5+ were pooled and used to isolate polysome-associated RNA.

**Supplemental Figure 2:**
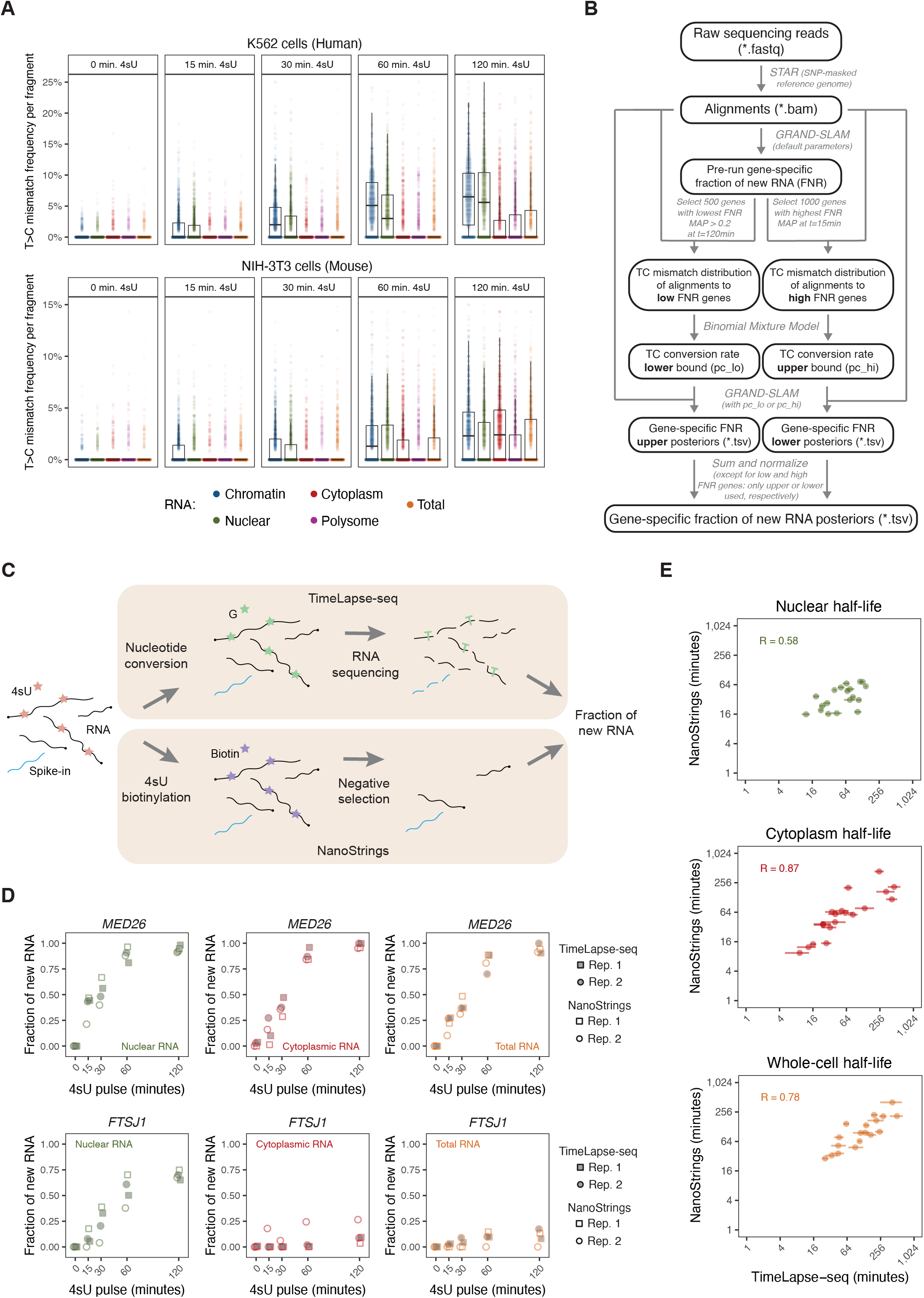
Nucleotide conversion analysis estimates the fraction of new RNA in a compartment- and time point-specific manner, related to Figure 1. (A) Frequency of T>C mismatches in RNA-seq reads relative to 4sU pulse durations. The frequency of mismatches is calculated for each read as the number of T>C mismatches over the total number of Ts per fragment (using both forward and reverse reads). A total of 1,000 reads are analyzed for each sample. Dots represent individual fragments. (B) Computational analysis pipeline for subcellular TimeLapse-seq data (see Methods for details). (C) Schematic for NanoStrings-based and subcellular TimeLapse-seq estimation of fraction of new RNA. Top: TimeLapse-seq analysis uses an oxidative nucleophilic-aromatic substitution reaction (Schofield et al., 2018) to recode the 4-thiouridine (4sU) molecules as cytosines, resulting in the incorporation of a guanine nucleotide during the reverse transcription step of library preparation. These are subsequently converted into cytosines during PCR amplification and identified computationally as T (genome) to C (sequencing read) mismatches during alignment. The fraction of new RNA per gene is estimated from sequencing reads as per (B). Bottom: NanoStrings-based analysis starts with the covalent biotinylationation of 4sU molecules, followed by the removal of 4sU-labeled RNAs by incubating the sample with streptavidin beads and retaining the supernatant (unbound RNAs). The number of RNAs per gene in the remaining sample (unlabeled RNAs) is determined by hybridization with NanoString probes. The fraction of 4sU-labeled is determined by normalizing the RNA counts to an unlabeled spike-in RNA and comparing to no 4sU control (see Methods for more detail). (D) Fraction of new RNA within nuclear (left), cytoplasmic (middle), and total RNA (right) compartments as measured by subcellular TimeLapse-seq and NanoStrings for two example genes (*MED26 and FTSJ1*) in human K562 cells. Two biological replicates (“rep”) are shown for each approach. (E) Correlation of fraction of new RNA between NanoStrings and subcellular TimeLapse-seq for all genes and compartments. The data shown in (D) is summarized by calculating a nuclear, cytoplasm, and whole-cell half-life for each gene included in the Nanostrings panel (n=20) with the fraction of new RNA from each technique. Half-lives are calculated from NanoStrings-based fraction of new RNA values with least squares estimates (see Methods for more detail). Each dot represents one gene. Mean half-lives between replicates for each technique are plotted with the Pearson correlation. The horizontal error bars represent the 95% credible intervals from the RNA kinetic flow model from subcellular TimeLapse-seq.

**Supplemental Figure 3:**
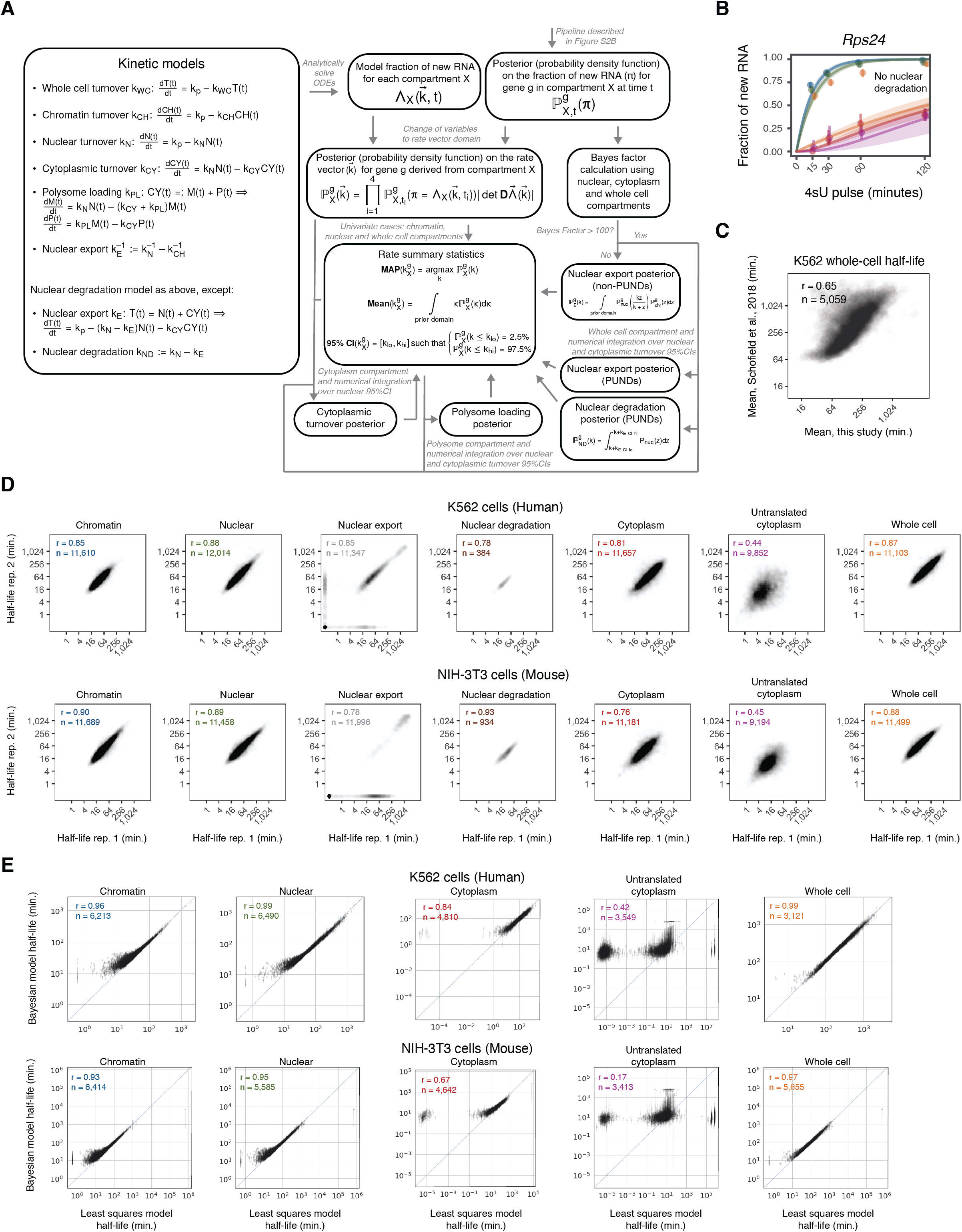
RNA flow can be modeled using a series of ordinary differential equations in a Bayesian framework, related to Figure 1. (A) Schematic of RNA kinetic modeling (see Methods for more detail). (B) *Rps24* model fits in the absence of nuclear degradation, related to Figure 1F (subcellular TimeLapse-seq in mouse NIH-3T3). Colors are consistent with Figure 1F. (C) Correlation between whole-cell (total RNA) half-lives measured in K562 cells in this study and in a previous study. The mean total RNA half-life between replicates, calculated using TimeLapse-seq following a single 4sU pulse (Schofield et al., 2018), was compared to the mean whole-cell half-lives between replicates in this study. Pearson’s correlation is shown and the number of genes is noted. Each dot represents one gene. (D) Reproducibility of RNA flow rates across biological replicates. For each flow rate, the mean half-life is compared between replicates. Pearson’s correlation is shown and the number of genes is noted. Each dot represents one gene. (E) Comparison between the Bayesian model and a least squares model. The Bayesian MAP half-life for each subcellular compartment for one replicate is compared with the least squares estimate. Pearson’s correlation is calculated and the number of genes is noted. Each dot represents one gene. Error bars indicate 95% credible interval of Bayesian half-lives.

**Supplemental Figure 4:**
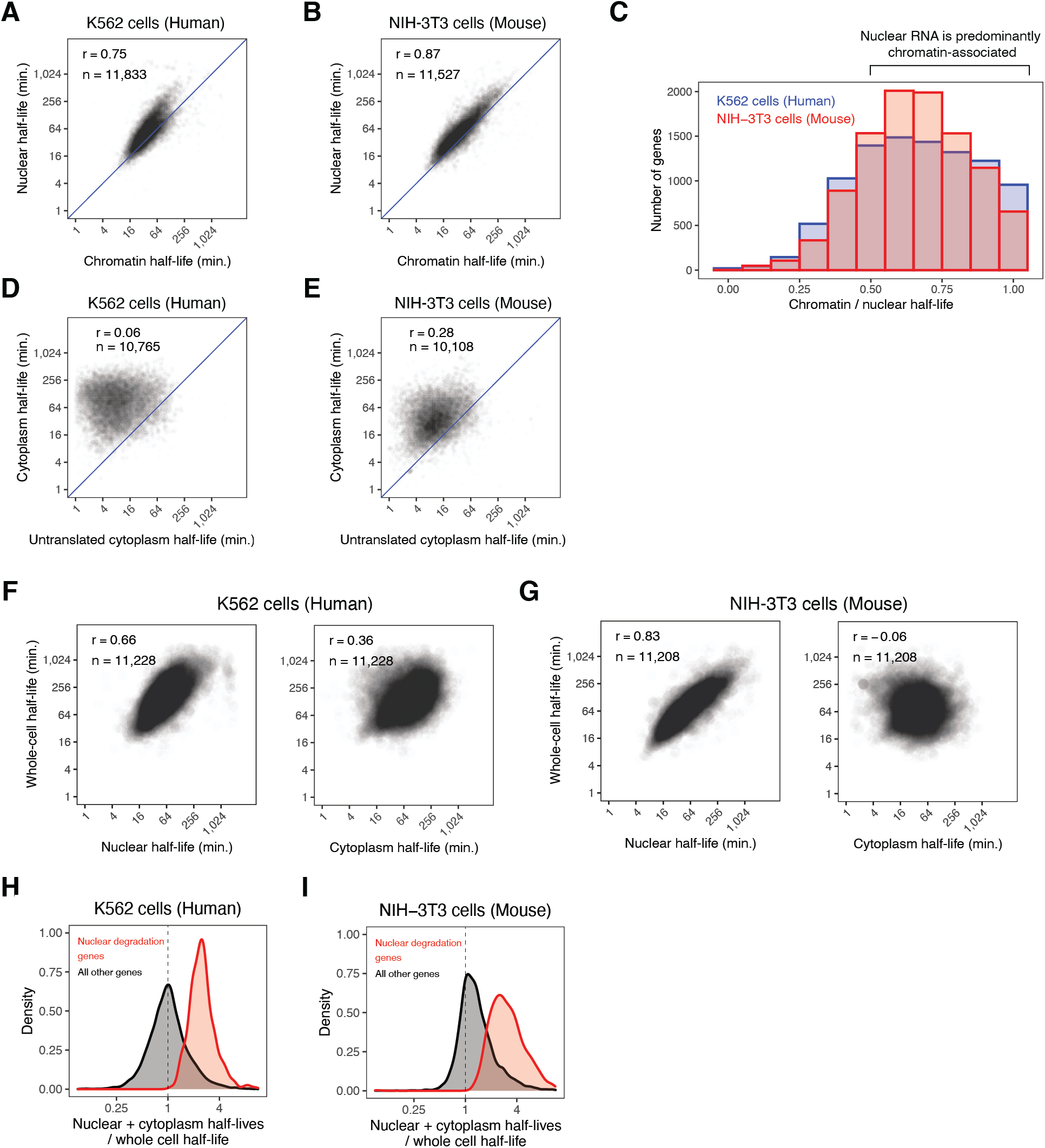
RNA flow rates show consistent genome-wide patterns between cell lines, related to Figure 1. (A) Correlation between chromatin and nuclear RNA half-lives in human K562 cells. Mean half-lives for each rate are compared with the Pearson correlation and number of genes noted. Each dot represents one gene. (B) Same as (A) in mouse NIH-3T3 cells. (C) Distribution of the ratio of chromatin to nuclear half-lives, showing that nuclear RNA is predominantly chromatin-associated (ratio>0.5) for a majority of genes. Data for both human K562 and mouse NIH-3T3 cells is shown as a histogram. (D) Correlation between untranslated cytoplasm and cytoplasm half-lives, showing that these rates are not related in human K562 cells. Mean half-lives of each rate are compared with the Peason correlation and number of genes noted. Each dot represents one gene. (E) Same as (D) in mouse NIH-3T3 cells. (F) Comparison of nuclear half-lives or cytoplasm half-lives to whole-cell half-lives in human K562 cells. Mean nuclear half-lives (left) or mean cytoplasmic half-lives (right) are compared to mean whole-cell half-lives with the Pearson correlation and number of genes shown. Each dot represents one gene. (G) Same as (H) in mouse NIH-3T3 cells. (H) Density distribution of the “predicted” whole-cell half-life (the sum of the nuclear and cytoplasm half-lives) divided by the observed whole-cell half-life. Genes with model fits without nuclear degradation are shown in gray and genes with model fits including nuclear degradation are shown in red. (I) Same as (H) in mouse NIH-3T3 cells.

**Supplemental Figure 5:**
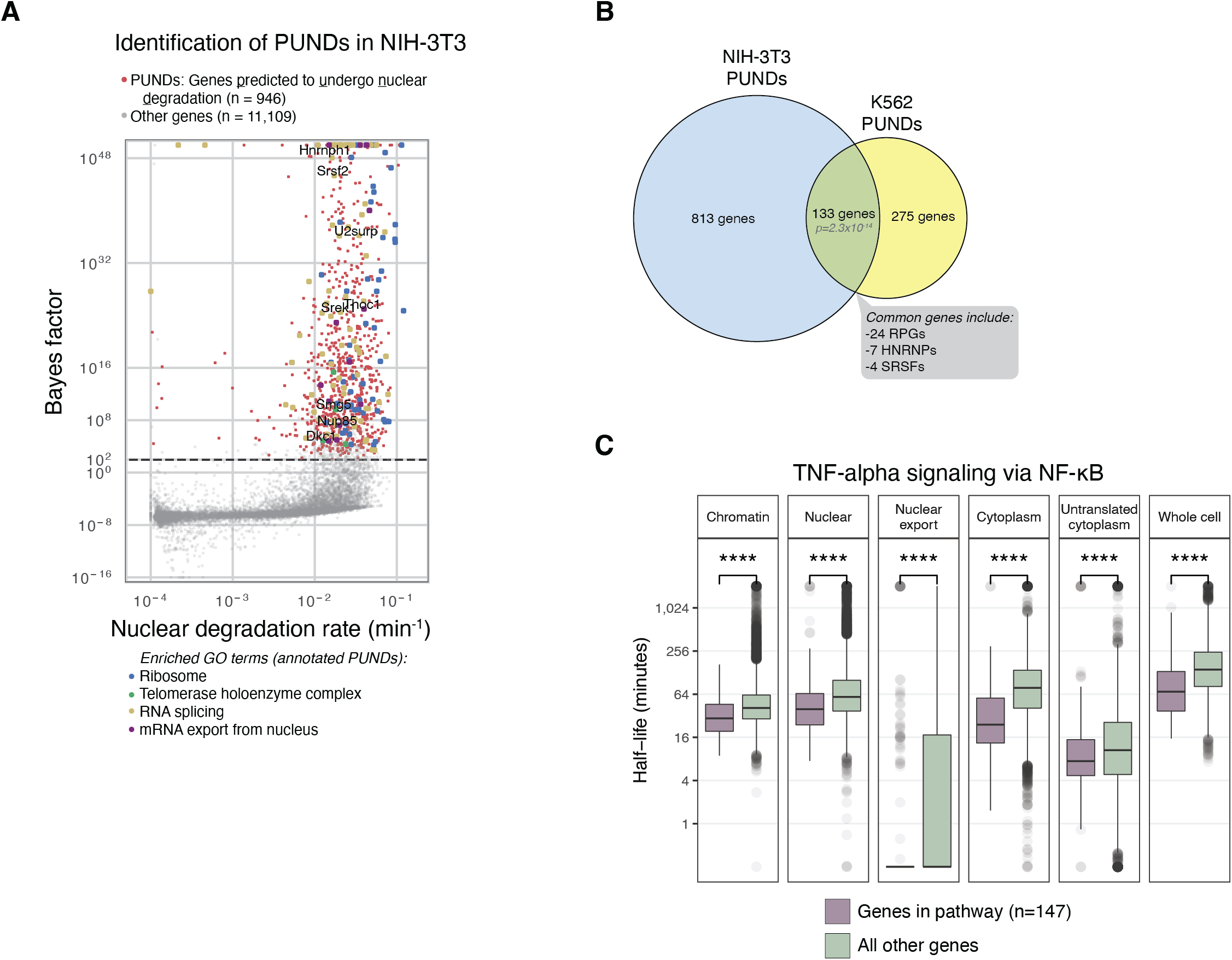
Functionally related genes exhibit similar RNA flow, related to Figure 2. (A) Identification of PUND genes in mouse NIH-3T3 cells. The Bayes factor and nuclear degradation rate are shown for each gene as in Figure 2A. (B) Venn diagram showing the number of unique and common PUND genes between mouse NIH-3T3 and human K562 cells. A Fisher’s exact test was performed to test the significance of the number of common genes. This common gene list included ribosomal protein genes (RPGs), heterogeneous nuclear ribonucleoproteins (hnRNPs), and SR splicing factors (SRSFs). (C) RNA flow rates for genes involved in TNF-alpha signaling via NF-κB, as defined by GSEA MSigDB (Subramanian et al., 2005). Subcellular half-lives were compared between groups using a Wilcoxon test (****: p<0.0001, ***: p<0.001, **: p<0.01).

**Supplemental Figure 6:**
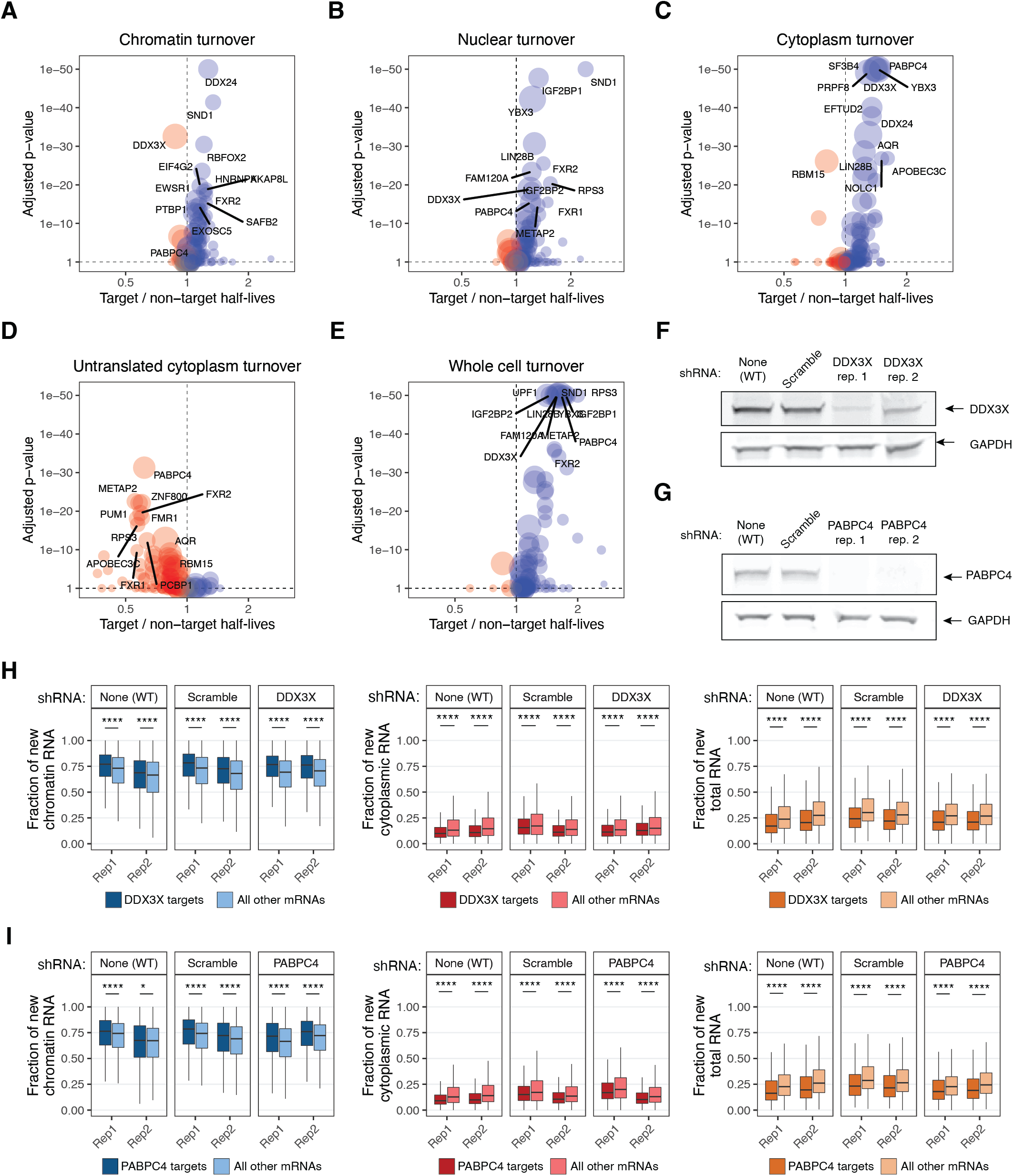
RNA binding proteins are associated with RNA flow rates, related to Figure 3. (A) Comparison of chromatin RNA half-lives between target and non-target mRNAs of each RBP according to Fig. 3A. The median chromatin half-life of the targets was compared to the median chromatin half-life of the non-targets with a Wilcoxon test (x-axis) with RBPs associated with slower target turnover in blue and RBPs with faster target turnover in red. The adjusted p-value following a Bonferroni correction for each RBP is indicated on the y-axis, and the size of the dot for each RBP indicates the number of target mRNAs. (B) Same as (A) for nuclear half-lives. (C) Same as (A) for cytoplasm half-lives. (D) Same as (A) for untranslated cytoplasm half-lives. (E) Same as (A) for whole-cell half-lives. (F) Confirmation of DDX3X protein knockdown in K562. Cells were transduced with lentivirus containing plasmids expressing a scrambled shRNA sequence or one targeting DDX3X and knockdown efficiency was monitored by western blotting in samples collected for subcellular TimeLapse-seq. Wild-type (non-transduced) cells were included as a control. (G) Same as (F) for PABPC4. (H) Fraction of new RNA in chromatin, cytoplasm, and whole-cell following DDX3X knockdown for target and non-target mRNAs. Fraction of new RNA MAP values were compared between targets and non-targets with a Wilcoxon test (****: p<0.0001, *: p<0.05). (I) Same as (H) for PABPC4.

**Supplemental Figure 7:**
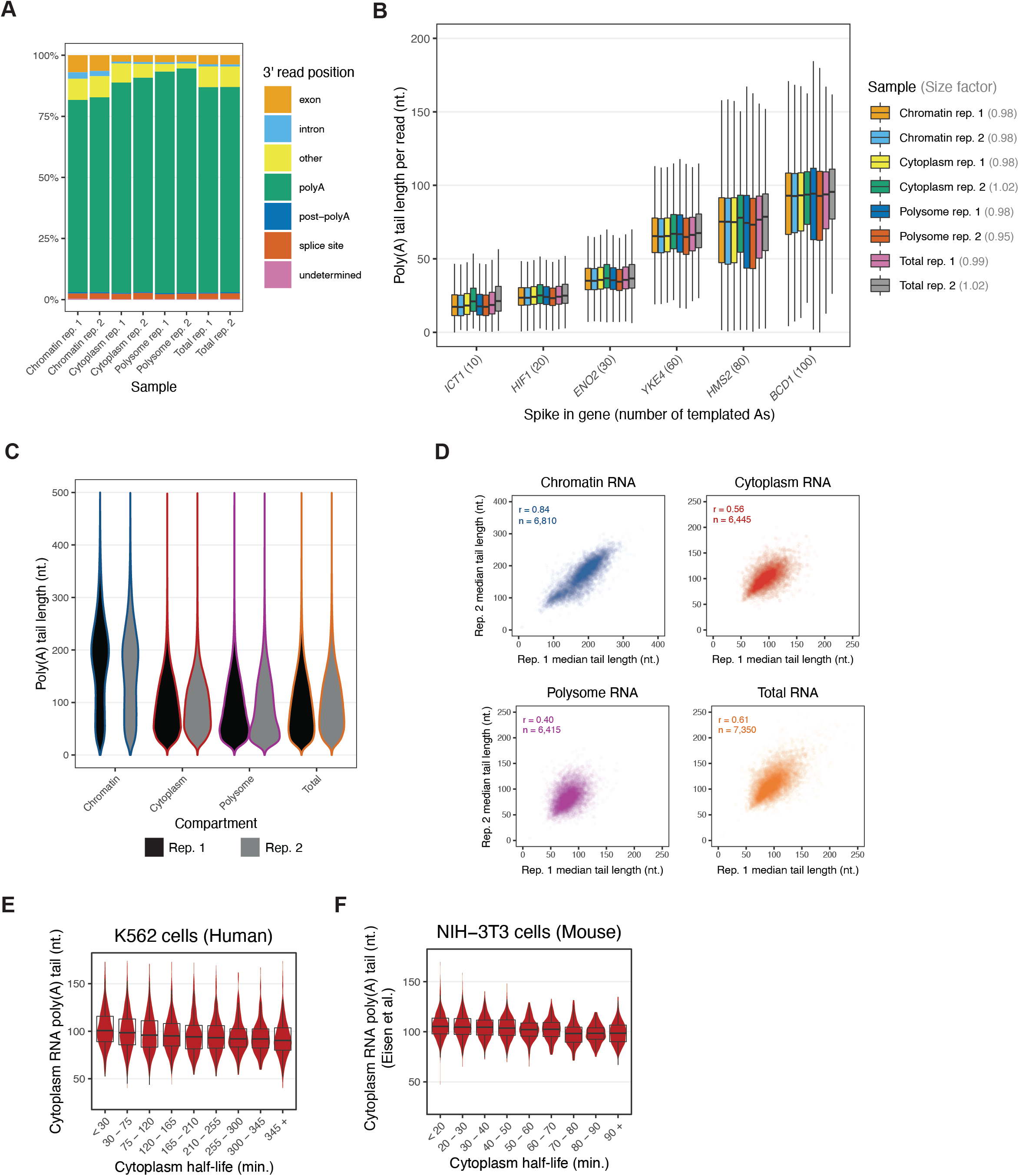
Poly(A) tail lengths are related to RNA flow rates, related to Figure 4. (A) Distribution of 3’ ends of poly(A)-selected RNA direct sequencing reads. The genomic region corresponding to the 3’ end of reads across all samples was determined according to (Drexler et al., 2020). (B) Distribution of poly(A) tail lengths for synthetic spike-in RNAs across nanopore sequencing runs. Six transcripts (shown on the x-axis) from *S. cerevisiae* with templated poly(A) tails ranging from 10 to 100 nucleotides were transcribed *in vitro* and added to each sample prior to nanopore library preparation. The median poly(A) tail length of each spike-in transcript within each sample was then used to calculate a poly(A) tail length size factor for each sample (noted in gray). Raw poly(A) tail lengths for each read were normalized to this size factor (see Methods for more details). (C) Distribution of poly(A) tail lengths per read, normalized to the synthetic spike-ins, across all samples and replicates. (D) Correlation of median compartment-specific poly(A) tail lengths between biological replicates. Median tail lengths are calculated for all genes containing >=10 reads for each compartment, with the Pearson correlation between biological replicates and the number of total genes noted. Each dot represents one gene. (E) Distribution of cytoplasm poly(A) tail lengths as a function of cytoplasm half-lives in human K562 cells. The median cytoplasm RNA poly(A) tail length is shown for all genes containing >=10 reads. (F) Distribution of cytoplasm poly(A) tail lengths as a function of cytoplasm half-lives in mouse NIH-3T3 cells. The mean steady state poly(A) tail length in cell line 1 (Eisen et al., 2020a) was compared to the cytoplasm half-lives (this study).

**Supplemental Figure 8:**
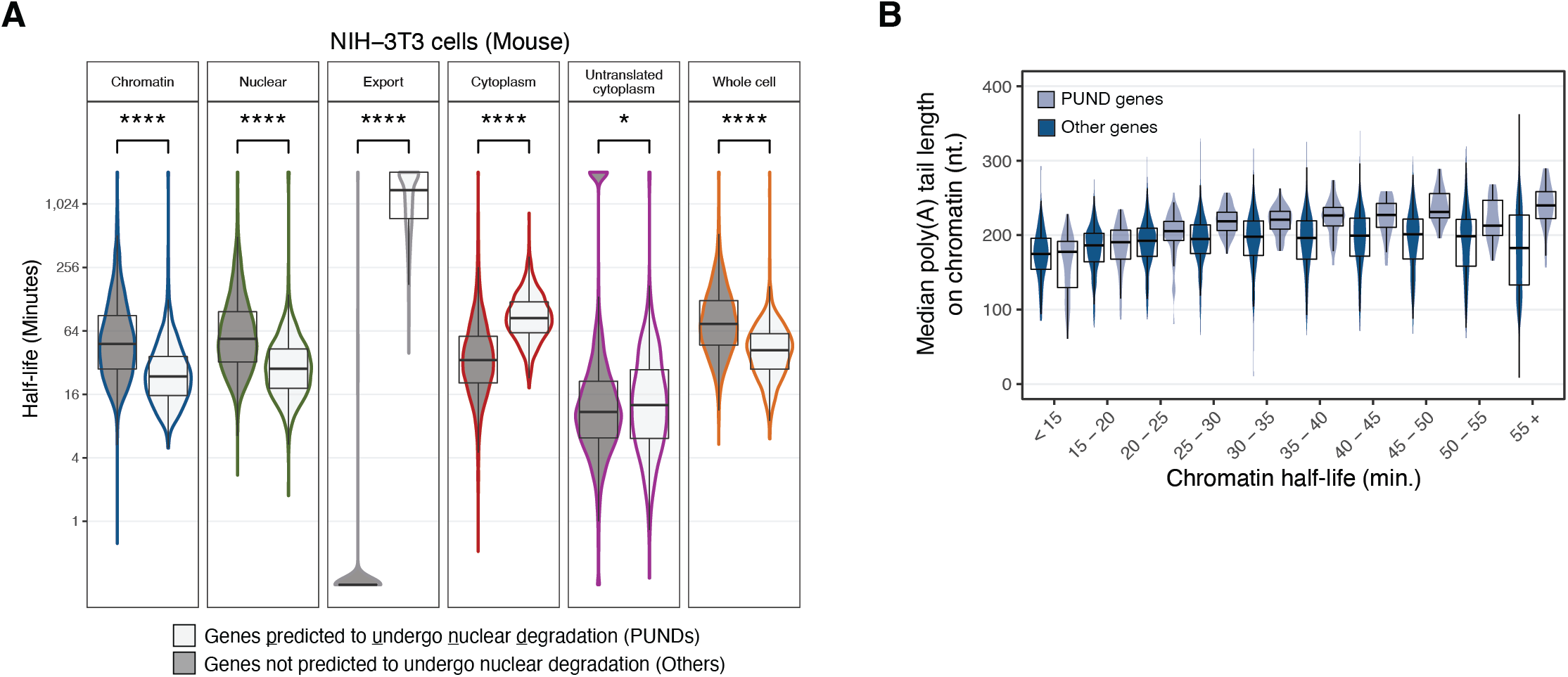
PUND phenotypes, related to Figure 5. (A) Comparison of half-lives of all PUND genes (n=946) and all other genes not predicted to undergo nuclear degradation across all RNA flow rates in NIH-3T3 using a Wilcoxon test (****: p<0.0001, *: p<0.05). (B) Distribution of median chromatin poly(A) tail lengths for PUND genes relative to all other genes as a function of chromatin half-life. The median poly(A) tail length was calculated for all genes containing >=10 reads. Within each bin, the lighter blue represents PUND genes while the darker blue represents other genes.

**Supplemental Figure 9:**
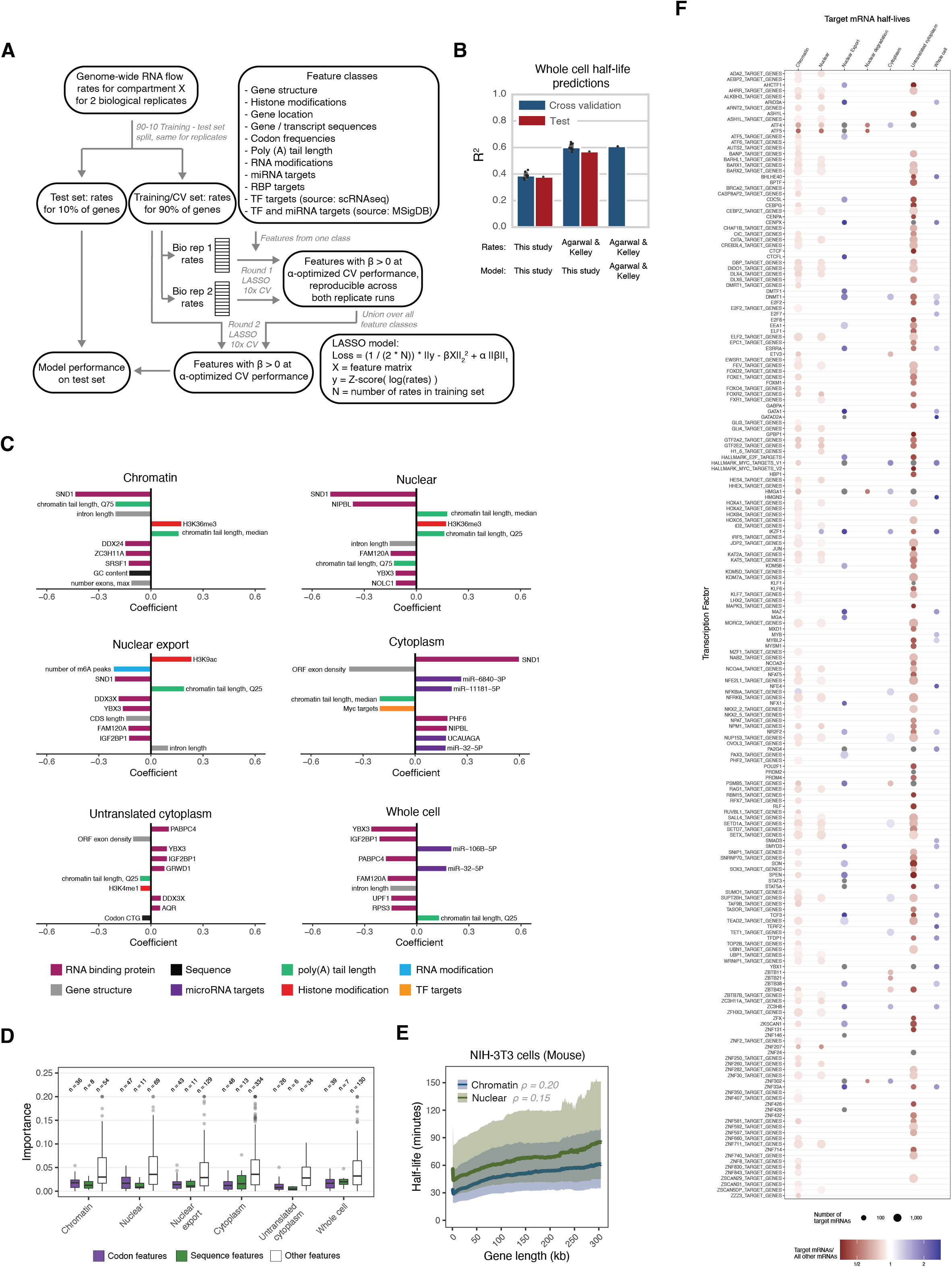
LASSO model predictions, related to Figure 6. (A) Schematic of LASSO feature selection and model training, 10x cross-validation and testing. (B) Comparison of our LASSO 10x cross-validation and test performance with alternative whole-cell turnover rate estimate and models. (C) The 10 individual features with highest importance for each RNA flow rate. Features are colored by family. (D) The importance of individual sequence and codon features compared to individual features of other families. (E) Continuous averages of chromatin and nuclear half-lives as a function of gene length in mouse NIH-3T3 cells (see Methods). Gene length was defined as the median genomic length of all transcripts per gene. Solid lines represent median half-lives and shaded ribbons represent the third quartile (top) and first quartile (bottom) of half-lives. (F) All transcription factors with targets that exhibited significantly fast or slow half-lives for target RNAs compared to non-target RNAs in both biological replicates (adjusted p<0.01, Wilcoxon test, Bonferroni multiple testing correction) across any RNA flow rate. The size of the dot indicates the number of target mRNAs with measured half-lives within each compartment and the color reflects the difference (red, faster; blue, slower) of the median target over non-target half-lives (see Methods for details).

